# Variation of human neural stem cells generating organizer states *in vitro* before committing to cortical excitatory or inhibitory neuronal fates

**DOI:** 10.1101/577544

**Authors:** Nicola Micali, Suel-Kee Kim, Marcelo Diaz-Bustamante, Genevieve Stein-O’Brien, Seungmae Seo, Joo-Heon Shin, Brian G. Rash, Shaojie Ma, Yanhong Wang, Nicolas A. Olivares, Jon Arellano, Kristen R. Maynard, Elana J. Fertig, Alan J. Cross, Roland Burli, Nicholas J. Brandon, Daniel R. Weinberger, Joshua G. Chenoweth, Daniel J. Hoeppner, Nenad Sestan, Pasko Rakic, Carlo Colantuoni, Ronald D. McKay

## Abstract

Better understanding the progression of neural stem cells (NSCs) in the developing cerebral cortex is important for modeling neurogenesis and defining the pathogenesis of neuropsychiatric disorders. Here we used RNA-sequencing, cell imaging and lineage tracing of mouse and human *in vitro* NSCs to model the generation of cortical neuronal fates. We show that conserved signaling mechanisms regulate the acute transition from proliferative NSCs to committed glutamatergic excitatory neurons. As human telencephalic NSCs developed from pluripotency *in vitro*, they first transitioned through organizer states that spatially pattern the cortex before generating glutamatergic precursor fates. NSCs derived from multiple human pluripotent lines varied in these early patterning states leading differentially to dorsal or ventral telencephalic fates. This work furthers systematic analysis of the earliest patterning events that generate the major neuronal trajectories of the human telencephalon.

## INTRODUCTION

Defining how cell types emerge in the forebrain is central to understanding the origins of normal and pathological function in the cerebral cortex (Geschwind and Rakic, 2013; Kwan et al., 2012b; Lein et al., 2017; Nowakowski et al., 2017; Sandberg et al., 2016; Wamsley and Fishell, 2017). The neocortex in mammals, including rodents and humans, is the product of fate transitions of radial glial cells (RGCs), which function as neural stem cells (NSCs), sequentially generating waves of post-mitotic neurons that migrate superficially from the ventricular germinal zones (VZ) to form the ontogenic columns of the cortical layers (Angevine and Sidman, 1961; Malatesta et al., 2000; Noctor et al., 2001; Rakic, 1974, 1988). This evidence has led to a sustained interest in defining how the commitment and transition from proliferative RGCs to excitatory cortical neuronal fate are controlled.

In the developing mammalian telencephalon, organizer centers secreting morphogenic signals emerge to pattern the cortical field prior to neuron specification (Geschwind and Rakic, 2013; Grove and Fukuchi-Shimogori, 2003; O’Leary et al., 2007; Sur and Rubenstein, 2005). Moreover, the excitatory and inhibitory neurons of the cortex emerge in two different zones, the dorsal and ventral telencephalon (Kwan et al., 2012b; Sandberg et al., 2016; Wonders and Anderson, 2006). In spite of its central importance, many features of this very early period when telencephalic regional identities are first acquired are not well understood, particularly in human. Recent reports of species-specific differences in corticogenesis are often focused on relatively late neurogenic stages when there is an enhanced genesis, in humans, of superficial neurons from the outer subventricular zone (oSVZ) (Namba and Huttner, 2017; Nowakowski et al., 2016; Zhu et al., 2018). However, the evolutionary expansion of the human cerebral primordium is evident from the earliest stages and is already prominent when RGCs produce the first glutamatergic neurons (Bystron et al., 2008; Geschwind and Rakic, 2013). Thus, there is a clear interest in defining how the early patterning mechanisms are coordinated to achieve discrete waves of neurogenesis.

Evidence of genetic risk for neuropsychiatric disorders has been found in the patterns of genes expressed in the neurogenic fetal cortex (de la Torre-Ubieta et al., 2018; Gulsuner et al., 2013; Parikshak et al., 2013; State and Sestan, 2012; Willsey et al., 2013; Xu et al., 2014). Moreover, risk associated genes have been identified in the *in vitro* functional phenotypes of NSCs derived from patient-specific induced pluripotent stem cells (iPSCs) (Brennand et al., 2015; Consortium, 2017; Fujimori et al., 2018; Lang et al., 2018; Madison et al., 2015; Marchetto et al., 2016; Mariani et al., 2015; Schafer et al., 2019). These studies defining the molecular and developmental origins of risk for brain disorders point to the importance of early telencephalic fate transitions in the onset of pathogenic mechanisms.

*In vitro* neural systems are central in modeling these early events in neurogenesis. The growth factors FGF2, insulin and other extracellular ligands, acting through the MAPK/ERK and PI3K/AKT pathways on the expression of cell cycle regulators, control the critical transition when proliferating cortical NSCs initiate neurogenesis, both during brain development and in cell culture (Adepoju et al., 2014; Androutsellis-Theotokis et al., 2006; Cattaneo and McKay, 1990; Johe et al., 1996; Lehtinen et al., 2011; Qi et al., 2017; Rash et al., 2011; Ravin et al., 2008; Vaccarino et al., 1999). Lineage analysis of rodent NSCs differentiating *in vitro* directly demonstrated rapid commitment of multipotent cells to neuronal or glial fates (Ravin et al., 2008). However, we still lack a comprehensive view of the molecular events regulating human NSC progression to post-mitotic cortical glutamatergic excitatory neurons.

Here, we modulated FGF2-MAPK signaling to control the developmental progression of mouse and human NSCs towards neurogenesis *in vitro*. We first strictly define the acute molecular events as NSCs commit to excitatory glutamatergic fates. Then, building on previous work (Edri et al., 2015; Sakaguchi et al., 2015), we show that when human NSCs derived from PSCs were serially passaged *in vitro*, they self-organize to transition through a sequence of dorsal telencephalic developmental stages. At early passages human NSCs expressed genes characteristic of the cortical hem, the dorso-caudal organizer zone of the telencephalon (Caronia-Brown et al., 2014), progressing later to cortical glutamatergic neuronal fates. We then derived multiple human iPSC lines that showed intrinsic variability in this early organizer state formation and in the consequent differentiation bias to dorsal or ventral telencephalic fates. This work shows that patterning mechanisms and commitment events that generate dorsal or ventral telencephalic neuronal fates are coordinated features that emerge under precise control in human NSCs *in vitro*. This 2-dimensional experimental system opens to further systematic analysis these early non-linear fate transitions that specify human cortical fates.

## RESULTS

### Waves of transcriptional dynamics during mouse *in vitro* neurogenesis are regulated by FGF2 signaling

To define the events of cortical neuron commitment we first used primary NSC cultures derived from mouse dorsal telencephalon at the beginning of neurogenesis, embryonic day 11.5 (E11.5). *In vitro*, rodent cortical NSCs proliferate maximally in presence of 10 ng/ml FGF2 and abruptly differentiate when this growth factor is withdrawn (Adepoju et al., 2014; Cattaneo and McKay, 1990; Ravin et al., 2008). Here, we modulated FGF2 signaling to assess the differentiation trajectory of NSCs. NSCs were exposed to a range of FGF2 doses (0.1, 1, or 10 ng/ml) for 48 hours after passage, and their later differentiation was monitored on day *in vitro* 15 (DIV 15) (Figure 1A). Consistent with a known role for FGF2 in antagonizing neurogenesis (Rash et al., 2011), the cultures exposed to lower (0.1 or 1.0 ng/ml) rather than the higher (10 ng/ml) dose of FGF2 generated more glutamatergic excitatory neurons of every subtype, as defined using established cortical layer specific markers (Kwan et al., 2012b; Molyneaux et al., 2007; Shen et al., 2006) (Figure S1A and B), and expressed significantly higher level of the pre- and post-synaptic proteins Synapsin 1 (SYN1) and Homer1 (HOM1) (Figure 1Fi and iii, S1C). Electrophysiological data also show regulation of the neuronal activity induced by early FGF2 treatment (Figure S1D). These results confirm that the FGF2 signaling status of NSCs exiting the cell cycle regulates the production and the functionality of glutamatergic neurons many days later. This *in vitro* experimental system will be further used here to explore cellular progression during neurogenesis, from proliferation to the post-mitotic state.

**Figure 1.**
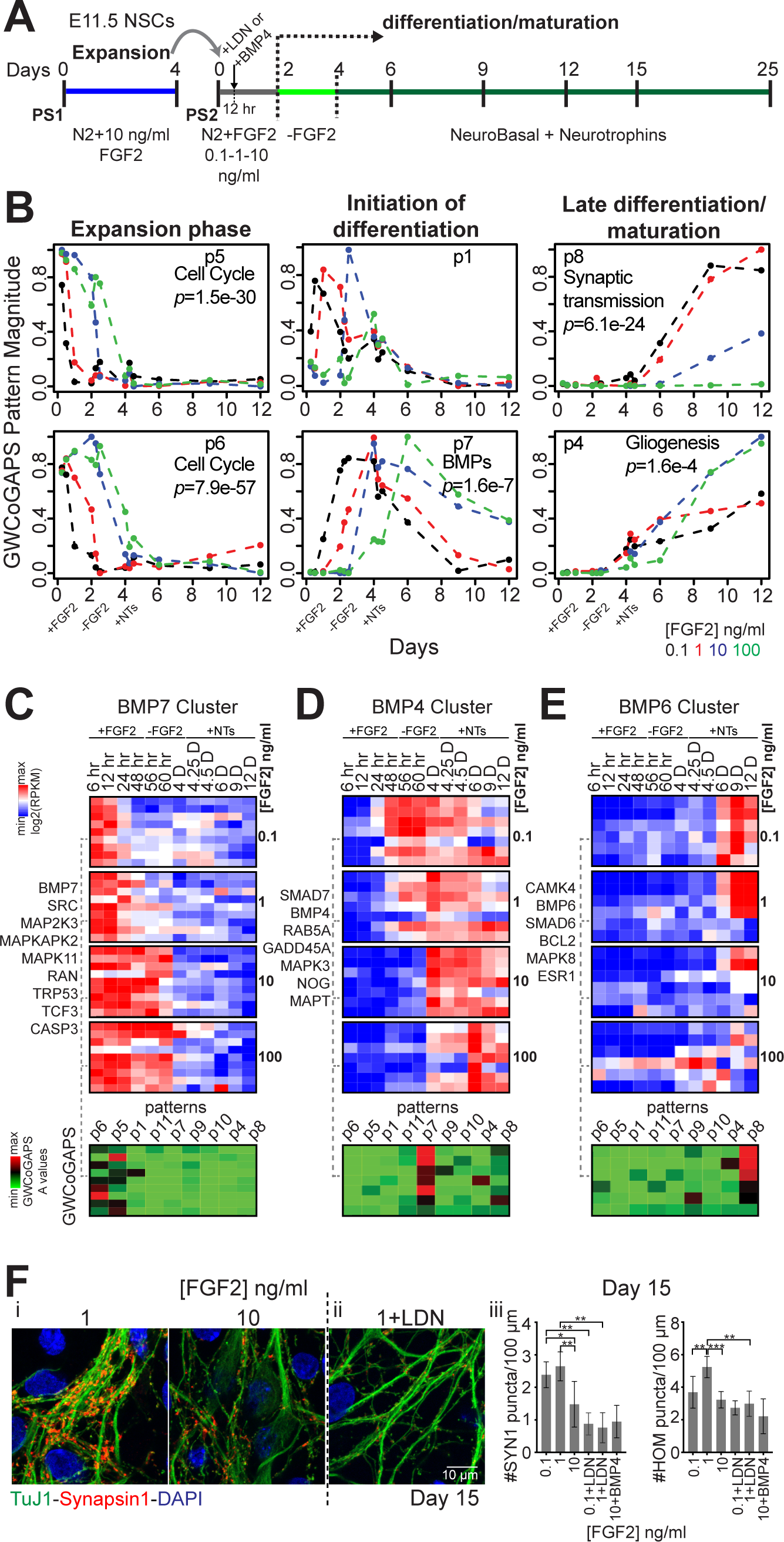
Transcriptional dynamics and BMP signaling across mouse *in vitro* neurogenesis (**A**) Experimental design. Passage 1 (PS1) NSCs were passaged into N2 + different FGF2 doses (PS2). Differentiation was induced by FGF2 withdrawal at DIV2. Neurotrophins (NTs) were added from DIV4. (**B**) 6 of 11 GWCoGAPS patterns shown in Figure S2. p-values indicate enrichment of GO categories in each pattern. (**C-E**) (Top) Expression dynamics of selected gene clusters annotated by “BMP receptor signaling” from NCI (full dendrogram in Figure S4), correlating with (C) BMP7, (D) BMP4 and (E) BMP6. BMP7 related genes were rapidly repressed in low FGF2. BMP4 related gene levels increased at the initiation of differentiation. BMP6 related genes were expressed at late time points. (Bottom) Gene weights in each GWCoGAPS pattern for the same upper genes. (**F**) Immuno-fluorescence images of SYN1 puncta in neurons cultured over astrocytes (i) at different FGF2 doses, or (ii) at 1 ng/ml FGF2 + LDN (100 nM). (iii) SYN1 and HOM1 puncta counts per 100 µm of neurite length for each condition, assessed by high-throughput image analysis. Mean ± St. Dev., t-test n > 5 fields per measurement. LDN or BMP4 were added 12 hours after NSC plating, and withdrawn with FGF2 at DIV2 (see A).

To gain insight into the transcriptional dynamics of NSCs progressing to excitatory neuron fates, mouse cortical NSCs were exposed to varying doses of FGF2 and total RNA was collected for RNA sequencing (RNA-seq), from 6 hours to 12 days of differentiation (Figure 1A). Principal component analysis (PCA), which identifies major axes of variation among the samples, defined progressive differentiation across the time in PC1 (Figure S2A), indicating that NSCs exposed to lower FGF2 concentrations moved more rapidly through this transition in gene expression. To explore these transcriptional dynamics at higher resolution, we employed the genome-wide CoGAPS (GWCoGAPS) non-negative matrix factorization (NMF) method. This method assigns weights for the contribution of every gene to a set number of patterns representing dominant changes in gene expression across all the *in vitro* samples (Fertig et al., 2014; Stein-O’Brien et al., 2017). GWCoGAPS analysis, identifying 11 patterns (p1-11), defined sequential waves of transcriptional extinction and induction as NSCs differentiated. Hierarchical clustering ordered these patterns into three groups associated with: (1) NSC expansion (p5 and p6); (2) initiation of neural differentiation (p1, p11, p7, p9 and p10); and (3) further maturation (p9, p10, p4 and p8) (Figure 1B, S2B and C). The early patterns (p5 and p6) were maintained in high FGF2 dose and enriched in cell cycle regulator genes, defined by gene ontology (GO) analysis. The initial FGF2 condition controlled sequential waves of gene expression in the first 6 days of differentiation (patterns p1, p11, p7 and p9). Importantly, GO analysis defined distinct end points for low and high FGF2 conditions that reflect either terminal neuro-(p8; low FGF2) or glio-(p4; high FGF2) genesis (Figure 1B). Hence, NSCs passaged into low FGF2 down-regulated cell cycle genes and traversed later steps of neuron differentiation more efficiently than in high FGF2 (Figure S3A-D). This high-resolution analysis defines dynamic waves of transcription initiated by variation in FGF2 signaling as NSCs exit the cell cycle, distinguishing differentiation trajectories that from the earliest times were biased towards either cortical neurons or glia.

### Early endogenous BMP signaling is required for mouse cortical neurogenesis

To address the events initiating differentiation of dorsal telencephalic neurons, we focused on BMP signaling that is known to oppose FGFs both in patterning the telencephalon *in vivo* and regulating NSC differentiation *in vitro* (Lehtinen et al., 2011; Lillien and Raphael, 2000; Mabie et al., 1999; Tiberi et al., 2012). When hierarchical clustering was used to analyze the dynamics of BMP signaling during *in vitro* neurogenesis, distinct waves of expression of BMP responsive genes correlated with particular BMP ligands were initiated by the early exposure to FGF2 (Figure 1B pattern p7 and C-E, S4A and B). Nuclear phosphorylated SMAD1/5 (pSMAD1/5) signal monitored a rapid transient induction of endogenous BMP in low FGF2 doses (Figure S4C), that paralleled the induction of mRNA expression for BMP4 and the BMP induced antagonist Noggin (Figure 1D and S4B).

To determine if this early BMP signaling had a significant effect on neurogenesis and neuronal maturation, we perturbed it 12 hours after plating NSCs, prior to BMP4 transcription (Figure 1A and S4B). The early inhibition of BMP signaling by the BMP type I receptor (BMPR1) inhibitor LDN193189, applied to NSCs exposed to low FGF2 dose, blocked both pSMAD1/5 induction at DIV 2 and the subsequent neurogenesis and synaptic maturation seen at DIV 15. Exposing NSCs to high FGF2 dose plus exogenous BMP4 boosted pSMAD1/5 levels but did not rescue the compromised synaptogenesis, suggesting that a specific BMP responsive cell state must be induced prior to the neuronal differentiation program (Figure 1F, S4D and E). We previously demonstrated that rodent telencephalic NSCs consist of a heterogeneous and dynamic population with rapidly varying lineage potential (Ravin et al., 2008). These data indicate that low FGF2 induces response to an endogenous and transient wave of BMP signaling that is required for cortical excitatory neuron commitment *in vitro*.

### Mouse cortical NSC subtypes show selective FGF2-induced BMP signaling activation and distinct fate bias

To further explore NSC diversity during cortical neurogenesis, we interrogated the surface expression of the tyrosine kinase receptors PDGFRα and EGFR which identify subsets of embryonic dorsal telencephalic NSCs generating neurons and glia *in vivo* and *in vitro* (Andrae et al., 2001; Lillien and Raphael, 2000; Park et al., 1999; Sun et al., 2005). Remarkably, the proportion of PDGFRα_high_ and EGFR_high_ cells was regulated by FGF2 dose in the first 2 days *in vitro*, and moreover, the expression of the receptors was mutually exclusive before decreasing at DIV 6 (Figure 2A and S5A). These data indicate that transient waves of EGFR and PDGFR positive cell states occurred during the differentiation progression of NSCs *in vitro*.

**Figure 2.**
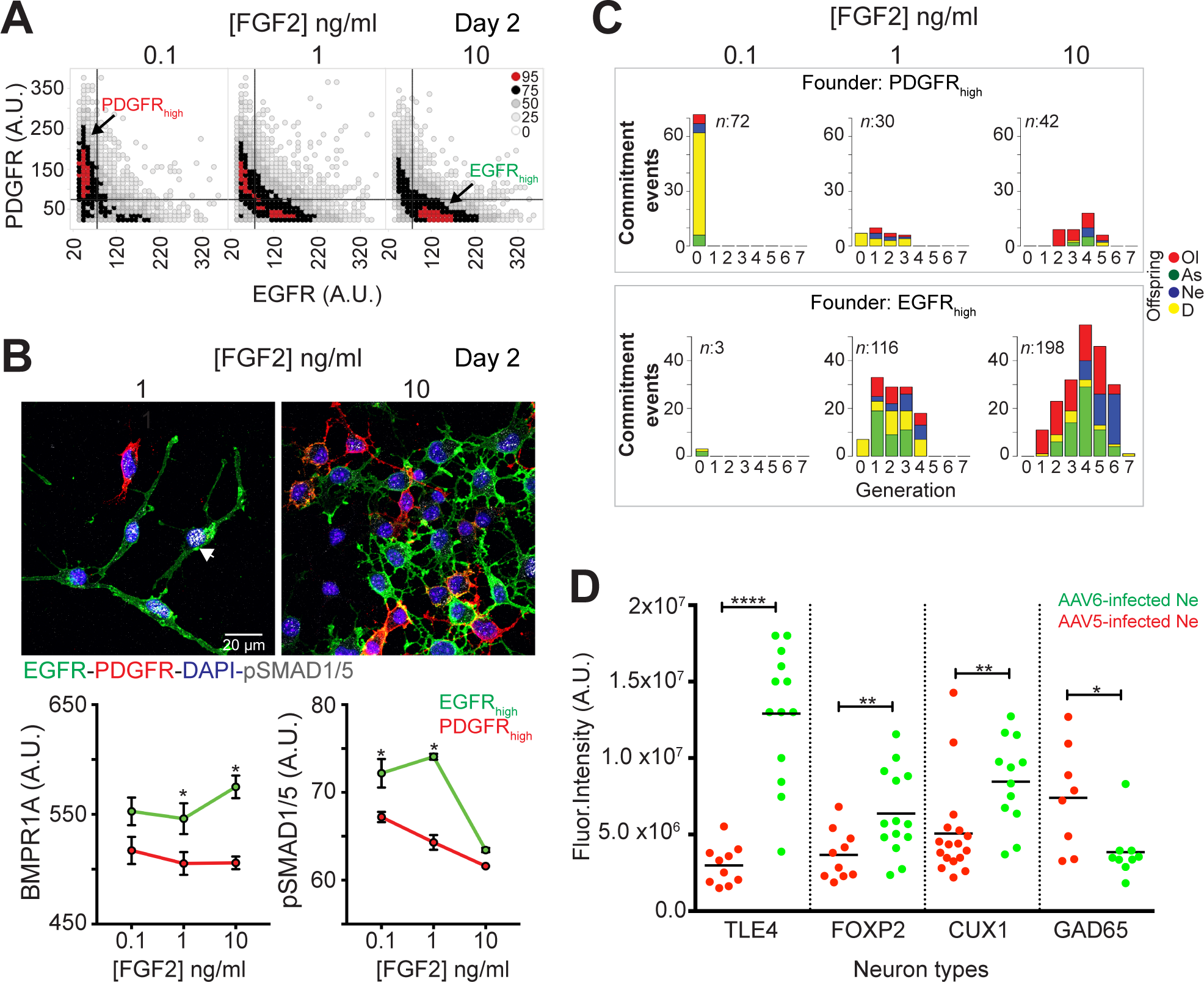
Mouse cortical NSC subtypes show selective BMP signaling activation and distinct fate bias. (**A**) Density plot of PDGFRα and EGFR expression in individually segmented cells from high-throughput image analysis (images in Figure S5A). Color key: percentile cell counts per bin. (**B**) (Top) Immuno-fluorescence images of pSMAD1/5 in EGFR_high_ and PDGFRα_high_ cells. Arrow shows pSMAD1/5 signals. (Bottom) Mean signal intensity of BMPR1A or pSMAD1/5 in EGFR_high_ and PDGFRα_high_ cells at DIV 2 from high-throughput image analysis, ± St. Dev. t-test. (**C**) Lineage analysis (see Video S1-3). Commitment events plot across cell generations from PDGFRα_high_ (top) or EGFR_high_ founder cells (bottom). An event is given by an initial progenitor generating offspring with same fate (see Experimental Procedures). Ol, oligodendrocytes; As, astrocytes; Ne, neurons; D, death (apoptosis). *n*= total commitment events. (**D**) Fluorescence intensity of each marker in individual AAV5-GFP- or AAV6-GFP-TuJ1^+^ neurons (Ne) at DIV 15. Mean values (lines). t-test.

Time-lapse recording and lineage analysis showed that FGF2 signaling controlled the cell cycle duration of EGFR_high_ cells in a dose dependent manner (Figure S5B), inducing also higher levels of phosphorylated FGF receptor (pFGFR) and pERK across all the doses (Figure S5C). However, no differences in FGFR2, FGFR1 and FGFR3 expression were seen between PDGFRα_high_ and EGFR_high_ cells (Figure S5C and not shown). These data suggest that FGF2 induced asymmetric signaling in the two subtypes. Higher expression of the neural fate transcriptional regulators HES1 and PAX6 was seen in EGFR_high_ cells, while the levels of the precursor marker SOX2 were higher in PDGFRα_high_ cells (Figure S5D). Extending the above data showing early induction of endogenous BMP signaling was required for cortical neurons to efficiently differentiate, higher BMPR1A expression and pSMAD1/5 signal was found in EGFR_high_ cells in low FGF2 (Figure 2B). These results indicate that BMP signaling was initiated in a transient EGFR_high_ NSC state.

The fate bias of these NSC subtypes was assessed by time-lapse recording to create lineage dendrograms linking EGFR_high_ or PDGFRα_high_ founder cells at DIV 1 to the derived neurons, oligodendrocytes and astrocytes, identified at DIV 6 by expression of TuJ1, O4 or GFAP (Figure S6Ai; Video S1-3). PDGFRα_high_ progenitors committed with similar proportions to oligodendrocytes or neurons in low FGF2, and predominantly to oligodendrocytes at the higher FGF2 dose. EGFR_high_ cells were tripotent, both in 1 and 10 ng/ml FGF2 (Figure 2C). Consistent with the transcriptomic data (Figure 1 and S3), the lineage analysis showed that faster cell cycle exit correlated with early neuron specification of the EGFR_high_ cells (Figure 2C and S6Aii). These results suggest that efficient neurogenesis was associated with an early wave of differentiating EGFR_high_ cells becoming acutely post-mitotic in low FGF2 conditions.

To lineage trace neurons at more mature stages of differentiation (DIV 15), we employed adeno-associated viruses (AAV) encoding green fluorescent protein (GFP) with preferential tropism for cells expressing either PDGFRα (AAV5) or EGFR (AAV6) (Di Pasquale et al., 2003; Jackson et al., 2006; Weller et al., 2010) (Figure S6B and C). AAV6-infected cells preferentially generated CUX1^+^, FOXP2^+^, or TLE4^+^ glutamatergic neurons, while AAV5-infected cells more GAD65 expressing putative GABAergic neurons with less complex morphologies (Figure 2D and S6D). These results define an EGFR_high_ BMP responsive cell state that efficiently produces cortical glutamatergic neuronal fates.

### Cortical neuron differentiation bias varies with the passage of human NSCs

Having defined the events of the transition to glutamatergic neurons from mouse cortical NSCs, we investigated neuronal commitment mechanisms in human dorsal telencephalic NSCs derived from PSCs *in vitro* (Edri et al., 2015; Mariani et al., 2012; Shi et al., 2012). Using standard neural induction protocols (Chambers et al., 2009; Edri et al., 2015), telencephalic NSCs were generated from the widely used human embryonic stem cell (ESC) line H9 that were then serially passaged in high FGF2 (Figure 3A). From passage (PS) 2 to 5, human NSCs were analyzed for the expression of fate regulators SOX2, HES1, HES5, OTX2, PAX6 and SOX21 (Figure S7A and B). The spontaneous emergence of the neural rosette-forming state at PS3 and PS4 and its subsequent decline provided a system to define the dynamics of neurogenic fate mechanisms in human.

**Figure 3.**
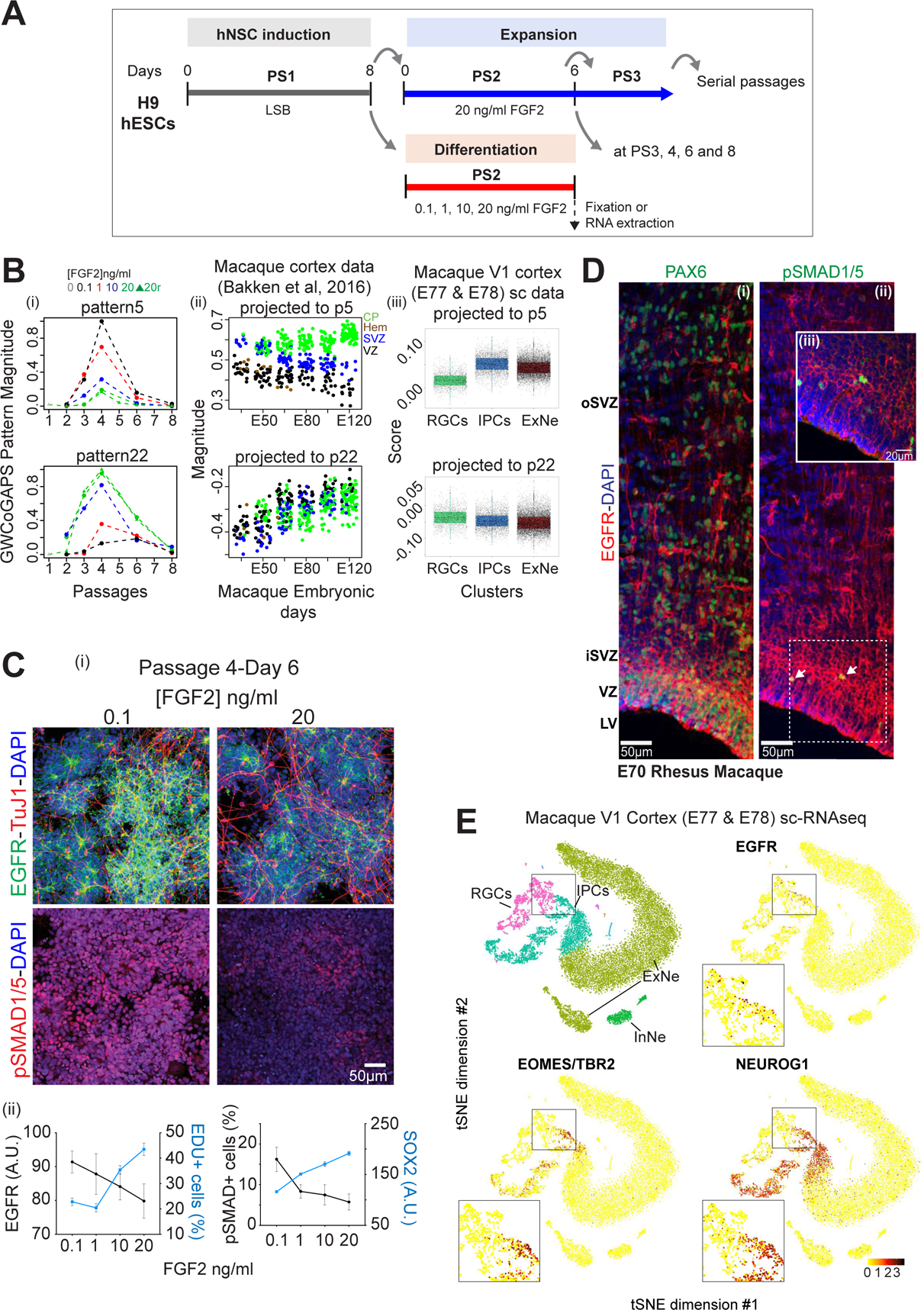
Cortical excitatory neuron fate bias at a specific passage of human NSCs (**A**) Scheme of H9 hESC differentiation into NSCs. N2 + LSB (LDN193189 + SB431542) was applied at passage 1, then hNSCs were serially passaged in N2 + 20 ng/ml FGF2. FGF2 modulation was applied at a specific passage for 6 days before RNA collection. (**B**) (i) GWCoGAPS patterns p5 and p22. FGF2 doses indicated. 0 refers to PS1; 20r are replicates for 20 ng/ml FGF2 (ii) Projections of macaque cortex microarray data from (Bakken et al., 2016). CP: cortical plate; Hem: cortical hem; SVZ: subventricular zone; VZ: ventricular zone. (iii) Projection of macaque V1 scRNA-seq data (**C**) (i) Immuno-fluorescence images of EGFR and TuJ1, or pSMAD1/5 expression. (ii) Mean fluorescence intensity of EGFR or SOX2 from high-throughput image analysis. Proportion of EDU^+^ or pSMAD1/5^+^ cells over total cells at DIV 6, for different FGF2 doses. Mean values ± St. Dev. **(D)** Immuno-histochemistry images of E70 macaque cortex sections for EGFR (red; all panels) with PAX6 (i), pSMAD1/5 (ii, arrows indicate some positive cells; iii, higher magnification). Nuclei stained with DAPI. oSVZ/iSVZ: outer/inner subventricular zone; VZ: ventricular zone; LV: lateral ventricle (**E)** t-SNE plots colored by annotated cells and indicated markers of macaque V1 scRNA-seq data. All major clusters are in Figure S9. EGFR^+^, TBR2^+^ and NEU-ROG1^+^ IPCs transitioning from RGCs are evident in the insets.

The neurogenic potential of the hNSCs was explored using FGF2 signaling modulation, as with the mouse system. Following serial expansion in high FGF2 from PS2 to 8, hNSCs from each passage were exposed to varying concentrations of FGF2 for 6 days before RNA collection for RNA-seq (Figure 3A). PC1 revealed transcriptional progression across passages that was independent of FGF2 dose (Figure S7Ci). Gene expression data can be visualized or “projected” into a low-dimensional space defined by another data set, allowing exploration of the transcriptional modules defined in one data set as they change in the other. To relate the dynamics of hNSCs *in vitro* to *in vivo* cortical development, we projected gene expression data from the developing human (Jaffe et al., 2018) or macaque neocortex (Bakken et al., 2016) into the transcriptional space defined by the individual gene weights from PC1, using projectR (Figure S7Cii and iii) (Stein-O’Brien et al., 2019); see Experimental Procedures). The projection analysis indicated the transcriptional dynamics identified by PC1 *in vitro* parallel human and macaque development as neurogenesis peaks *in vivo*.

To characterize these transcriptional dynamics at a more granular level, we explored a set of 24 patterns (p1-24) defined by the GWCoGAPS algorithm in this *in vitro* hNSC RNA-seq data (Figure S8A). Three of these patterns (p5, p6, p22) showed transcriptional differences across FGF2 dose that peaked at PS4, when neural rosettes were most abundant (Figure 3Bi and S8A). Projection of the developing macaque cortex gene expression data (Bakken et al., 2016) into these patterns shows that p5 specifically identifies a gene signature more highly expressed in the forming cortical plate (CP) than in the germinal SVZ and VZ domains throughout this neurogenic period (Figure 3Bii and S8B).

To more precisely relate these *in vitro* transcriptional dynamics to discrete cell types of the developing primate neocortex, we generated a single-cell mRNA sequencing (scRNA-seq) data set from 2 macaque fetal visual cortex (V1) samples collected at E77 and E78, using 10x Genomic platform. Data from 17161 single cells passed quality control measures and were included in the present study. Using unsupervised clustering, we identified major cell clusters including RGCs, intermediate neuronal precursor cells (IPCs), excitatory neurons (ExNe), interneurons (InNe) (Figure S9A and B). Projection of this monkey fetal V1 or human developing cortex scRNA-seq data (Nowakowski et al., 2017) into the hNSC GWCoGAPS patterns confirmed that p5 correlates with post-mitotic IPC and excitatory neuron signatures *in vivo* (Figure 3Biii and S9Ci-iv). Pattern p22 induced by high FGF2 was not associated with an excitatory neurogenic signature in either projection (Figure 3B and S8B). These results indicate that the *in vitro* appearance of new-born cortical neurons was favored specifically at PS4 by acute reduction of FGF2 and, importantly, that gene expression dynamics occurring during *in vitro* neurogenesis parallel those in the neurogenic primate neocortex.

Immuno-fluorescence was used to further explore the differentiation of PS4 hNSCs into post-mitotic neurons. The data show that low FGF2 favored a switch from proliferating (EDU^+^), SOX2^+^ hNSCs to SOX2^-^, PAX6^-^, TuJ1^+^, EGFR_high_, DCX^+^ post-mitotic (EDU^-^) new-born neurons (Figure 3C, S8Ciii-v and D). Elevated pSMAD1/5 immunoreactivity showed that this transition from dividing precursors to young neuroblasts induced by low FGF2 was associated with a burst of endogenous BMP signaling occurring most efficiently at passage PS4 (Figure 3C and S8Cvi). These results suggest that post-mitotic neurons with cortical glutamatergic features were specified by a conserved mechanism in human and mouse NSCs and, importantly, that hNSCs differ with passage in the generation of pertinent intermediates as excitatory cortical neuronal fates were specified (Figure S8Ci-iv).

### EGFR-BMP signaling interaction defines a neurogenic transition state in RGCs of the developing primate cortex

Next, we investigated whether the progression of the NSC states seen *in vitro* recapitulates fundamental events of RGC development *in vivo.* In primates, the expression of PAX6 defines the neurogenic potential of RGCs in the VZ and marks committed neurons in the oSVZ (Hansen et al., 2010; Mo and Zecevic, 2008). In the E70 macaque dorsal parietal cortex we detected EGFR and PAX6 co-expression in RGCs of the VZ/iSVZ and in committed neurons delaminating through the oSVZ (Figure 3Di and ii, S9Di). The induction of nuclear pSMAD1/5 detected in EGFR_high_ expressing RGCs indicates BMP signaling response, suggesting transition of these cells to neurogenic precursors (Lehtinen et al., 2011; Li et al., 1998; Saxena et al., 2018) (Figure 3Dii and iii). Moreover, we found EGFR_high_ precursors expressing the neurogenic transcription factor EOMES/TBR2 in iSVZ, confirming neuronal commitment of these cells (Figure S9Dii and iii) (Arnold et al., 2008; Englund et al., 2005; Pollen et al., 2015). Interestingly, EGFR staining was not detected in CP cells expressing later neuronal markers such as TBR1 (Figure S9Di, ii and iv).

We explored this transition further in the monkey fetal V1 scRNA-seq data and identified a population of early neuronal IPCs progressing from RGCs, expressing EGFR, TBR2, SOX2, NEUROG1 (Figure 3E and S9B). These results are consistent with the hypothesis that hNSCs *in vitro* model a mid-neurogenic phase of primate cortex development, when an EGFR_high_ expressing RGC population employs BMP signaling in a transition state from proliferative cells to committed glutamergic cortical neurons.

### Early passage human NSCs show cortical organizer identities

To understand the origins of the neurogenic peak, we focused on NSC states prior to PS4. Five GWCoGAPS patterns from the *in vitro* hNSC data showed the highest gene expression at PS2 and PS3 (p8, p11, p2, p4, p19) (Figure 4Ai and S10Ai). Of these, pattern p2, 11, 4 and 19 but not p8 were dependent on the initial FGF2 status of the cells. These transcriptional dynamics suggest these early stages of hNSCs also undergo an FGF2 regulated developmental progression that we explore further here.

Projection of the developing macaque cortex gene expression data (Bakken et al., 2016) into these early passage hNSC GWCoGAPS patterns showed that p8 and the low FGF2 patterns p11 and p2 expressed genes characteristic of the cortical hem domain (Figure 4Aii and S10B). Importantly, neither the early high FGF2 patterns p4 and p19, nor the later patterns p5 and p22 (described above), showed enrichment of genes of the cortical hem domain (Figure S10Aii and 3Bii). This analysis indicates that low FGF2 conditions at early passages induced elements of the hem transcriptional signature. The cortical hem is a source of BMP and WNT morphogens that pattern the dorso-caudal domain of the telencephalon (Caronia-Brown et al., 2014; Grove et al., 1998). In order to monitor the emergence of this cellular organizer identity across the hNSC passages, we used the gene expression data from the developing macaque cortex (Bakken et al., 2016) to generate a list of hem-specific genes of the primate telencephalon (Figure 4B; see Experimental Procedures). Passages 2 and 3 were enriched in these genes including LMX1A, WNT8B, WNT3A, BMP2/4/6/7, RSPO1/2/3 and BAMBI indicating coordinated transient expression of the cortical hem transcriptional signature, sensitive to FGF2 dose, at early passages of *in vitro* hNSCs (Figure 4B). The expression patterns of genes such as FGF8, TTR and SFRP2 at these early passages suggest that other organizer states including rostral patterning center (RPC), choroid plexus and antihem were also represented (Assimacopoulos et al., 2003; Sakaguchi et al., 2015; Storm et al., 2006) (Figure S10C).

**Figure 4.**
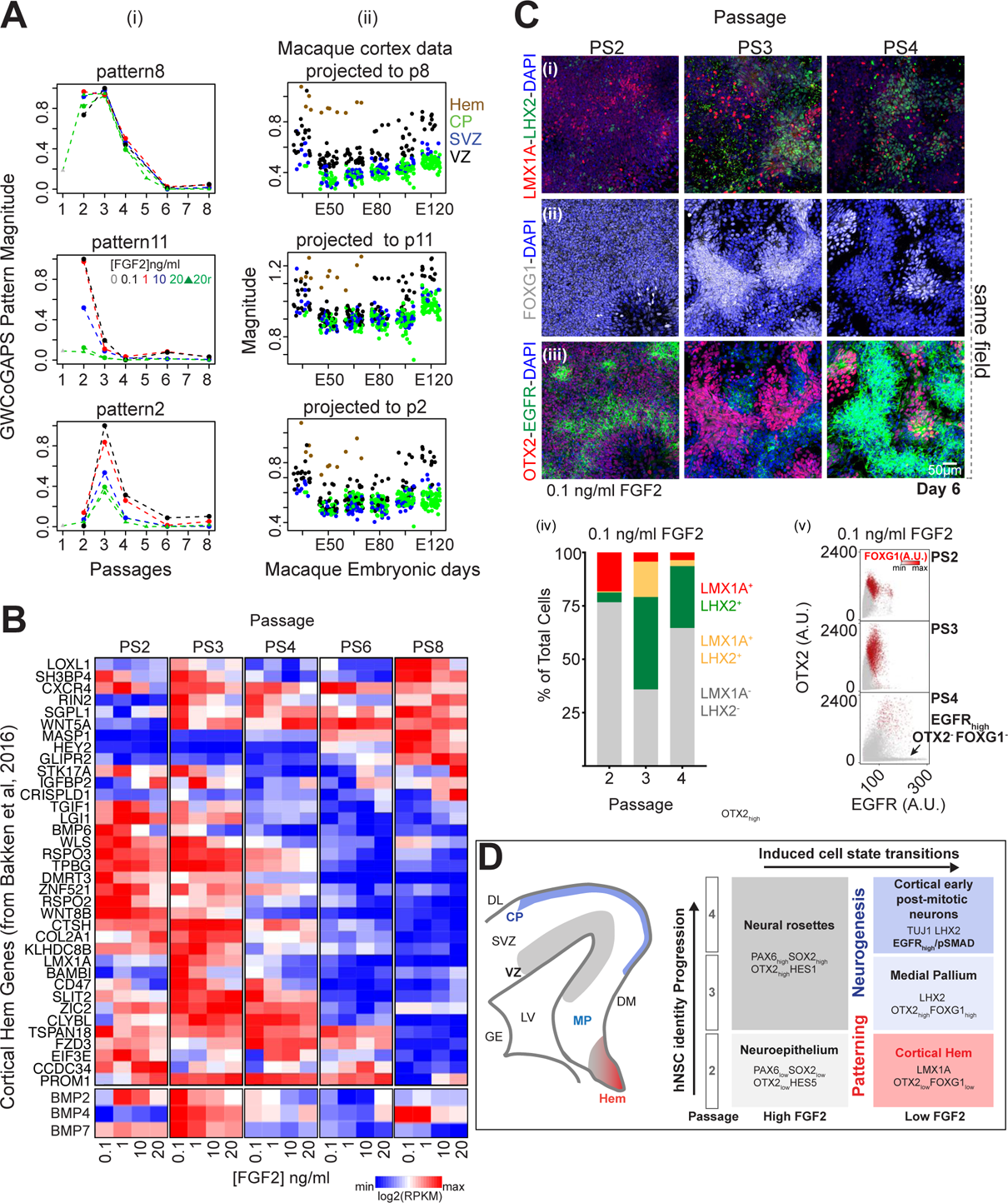
Cortical organizer identities of early human NSCs before progressing to neuronal fates. (**A**) (i) GWCoGAPS patterns p8, p11 and p2. FGF2 doses indicated. 0 refers to PS1; 20r are replicates for 20 ng/ml FGF2 (ii) Projections of macaque cortex data from (Bakken et al., 2016). (**B**) Expression of hem related genes. Gene list derived from panel (Aii) contrasting hem with other cortical regions from (Bakken et al., 2016). BMP2, 4 and 7, not derived from the gene list of Aii, are in separate heatmap at bottom. (**C**) Immuno-fluorescence images of (i) LMX1A and LHX2, (ii) FOXG1, (iii) EGFR and OTX2, in cells cultured with 0.1 ng/ml FGF2 for 6 days from passage 2 to 4. FOXG1 (ii) and EGFR/OTX2 (iii) are from same field. (iv-v) From high-throughput image analysis: (iv) proportion of LMX1A and LHX2; (v) scatter plot of EGFR vs OTX2 expressing cells, colored by FOXG1 level. **(D)** Model of the hNSC state progression *in vitro*. (Left) View of a coronal section of the developing mammalian telencephalon. CP: cortical plate; SVZ: subventricular zone; VZ: ventricular zone; MP: medial pallium; GE: ganglionic eminence; LV: lateral ventricle. DM/DL: dorso-medial/-lateral. (Right) Distinct states of *in vitro* hNSCs indicated by markers. State transition of hNSCs induced by low FGF2 showing lineage progression from hem state at passage 2 to cortical neurogenic identity at passage 4. The transition from NSCs to early post-mitotic neurons is defined by EGFR_high_ cells responsive to BMP signaling.

Consistent with the known function of LMX1A in the specification of neural organizers including the cortical hem in the mouse (Chizhikov et al., 2010), the expression of this transcription factor was highest at PS2 in low FGF2 and subsequently decreased (Figure 4Ci and iv). In contrast, FOXG1 and LHX2, which are required in the mouse to form the medial pallium and lateral cortex and to repress the formation of the cortical hem (Bulchand et al., 2001; Hanashima et al., 2007; Mangale et al., 2008; Molyneaux et al., 2007; Monuki et al., 2001), were expressed more prominently at later passages (Figure 4Ci, ii, iv and v). At PS3, the majority of cells expressed LMX1A or LHX2, suggesting co-emergence of the caudal hem organizer and precursor cells of the cortical field which then become dominant in PS4 (Figure 4Civ). The progressive appearance of cells co-expressing OTX2, a transcriptional regulator of the choroid plexus and cortical hem (Sakaguchi et al., 2015), and FOXG1 at PS3, followed by OTX2^-^ FOXG1^-^ EGFR_high_ cells at PS4 (Figure 4Cii, iii and v), is consistent with the progression of hNSCs from a hem to cortical neurogenic identity (Figure 4C and D). This coordinated change indicates that cortical patterning and dorsal excitatory neuronal specification mechanisms are efficiently executed across early passages of human neural precursors generated *in vitro*.

### Human NSC line variation in organizer states results in divergent neuronal fate trajectories

To address the stability of these early telencephalic differentiation steps we analyzed multiple human iPSC lines. 6 hiPSC lines [2063-1, −2; 2053-2, −6; 2075-1, −3] were generated from scalp fibroblasts of 3 donors and differentiated towards forebrain fates. Aiming to probe intrinsic developmental bias in early steps of telencephalic commitment in different human iPSC lines, we employed a 32-day differentiation protocol including the DKK1 mimetic XAV939 (X), which inhibits BMP and WNT signaling promoting anterior fates (Glinka et al., 1998). To limit FGF2 mediated effects on regional identity of the NSCs and neurogenesis (Hendrickx et al., 2009; Korada et al., 2002; Raballo et al., 2000; Rash et al., 2013), the neuronal differentiating conditions X + LSB (XLSB) were applied directly to the iPSCs, without intervening passages of the NSCs in presence of FGF2 (Figure 5A).

**Figure 5.**
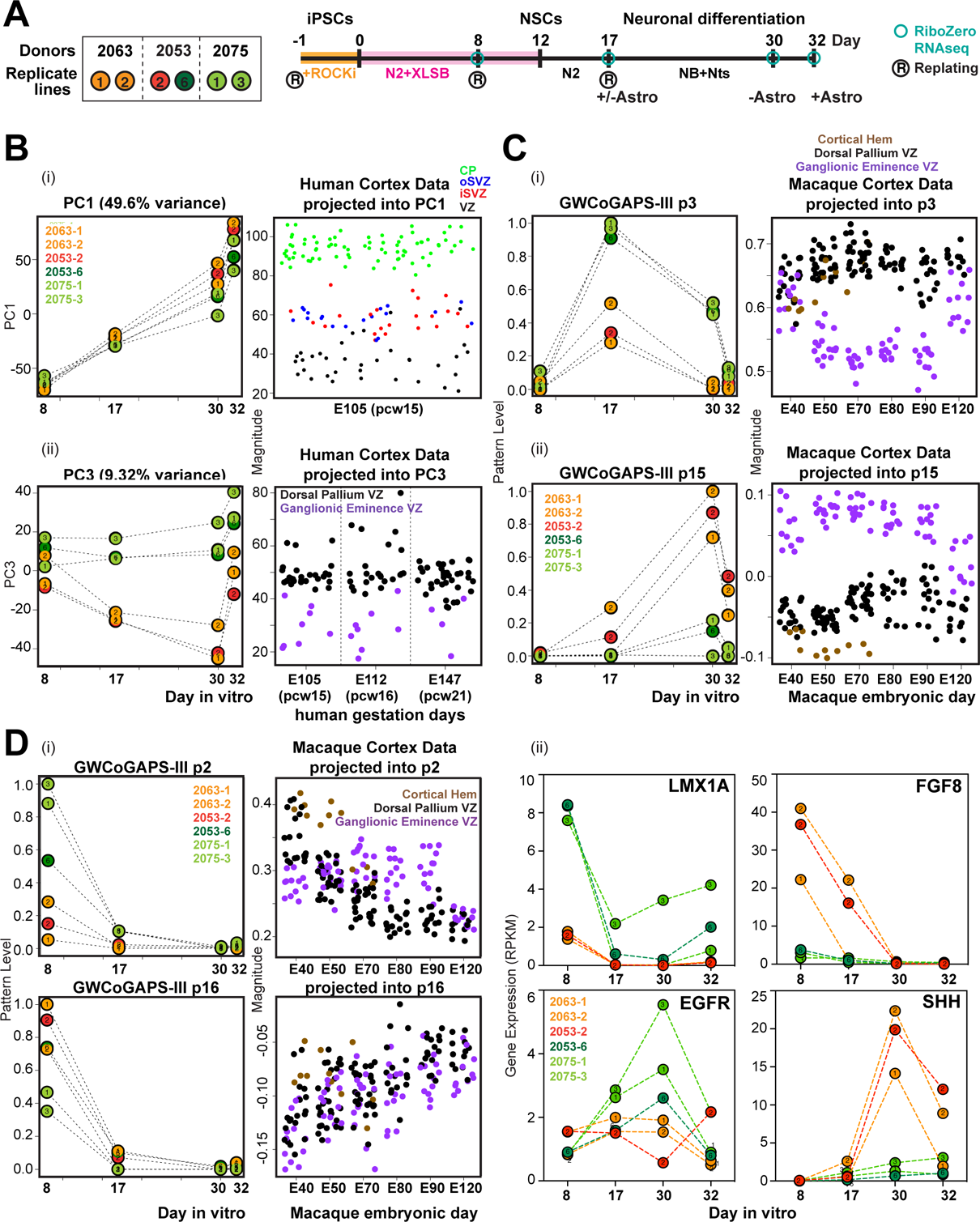
Human NSC line variation in organizer states results in divergent neuronal fate trajectories. (**A**) (Left) 6 hiPSC lines from 3 donors. Colors indicate divergent neuronal trajectory bias (see below). (Right) Experimental design. hiPSC lines were passaged in mTesR + Rock inhibitor then differentiated into NSCs in N2-B27 + XLSB medium for 12 days (Maroof et al., 2013). Sample collection and passages indicated. On day 17, NSCs were passaged and terminally differentiated with or without astrocytes. RNA was collected on day 30 (NSCs - Astro) or 32 (NSCs + Astro). (**B**) Distinct forebrain trajectories within 6 lines. (Left) Progression of neural differentiation represented by PC1 (i) and emergence of divergent trajectories represented by PC3 (ii). (Right) Projection of human cortex data (Miller et al., 2014). CP: cortical plate; oSVZ/iSVZ: outer/inner subventricular zone; VZ: ventricular zone. (**C**) (Left) Dorsal or ventral trajectories revealed by GWCoGAPS-III p3 (i) and p15 (ii). (Right) Projection of macaque cortex data (Bakken et al., 2016). (**D**) Hem genes are highly expressed in lines with dorsal lineage bias. (i) (Left) Divergent bias of lines at day 8 revealed by GWCoGAPS-III p2 and p16. (Right) Projection of macaque cortex data. (ii) Expression of the indicated genes across neural differentiation of the 6 hiPSC lines.

To analyze the differentiation states induced by this treatment, RNA was collected at different time points for sequencing analysis. PC1 of the gene expression data showed that all the samples progressed with similar kinetics through a common landscape of differentiation, promoted with similar efficiency in all 6 lines by growing NSCs either with or without rat astrocytes, known to facilitate neuronal maturation (Figure 5Bi). NSC fate regulators (SOX21, OTX2 and HES3) had high expression at early time points, in contrast to regulators of neuronal function (NEFL, SYP, SYT4, and SNAP25) expressed at later times (Figure S11A). Projection of gene expression data from micro-dissected regions of fetal human cortex (Miller et al., 2014) into PC1 demonstrated that the *in vivo* spatio-temporal progression that generates post-mitotic neurons of the CP maps onto the time course delineated in this PC1 (Figure 5Bi).

In contrast to PC1 where all 6 iPSC lines progressed equivalently, PC3 identified distinct patterns of gene expression on days 17 and 30 in two groups of cell lines: [2063-1, −2 and 2053-2] and [2053-6 and 2075-1, −3] (Figure 5Bii). Projection of the fetal human cortex gene expression data (Miller et al., 2014) into PC3 distinguished cell lines with differential expression of dorsal pallium [2053-6 and 2075-1, −3] or ventral ganglionic eminence genes [2063-1, −2 and 2053-2] (Figure 5Bii). Dorsal telencephalic regulators such as PAX6, FEZF2, NEUROD4 and NEUROG2 were highly expressed in one group of cells (2053-6, 2075-1 and −3), while regulators of ventral telencephalic fates such as NKX2-1, FOXG1, LHX6/8, SHH and DLX genes (Sandberg et al., 2016) were highly expressed in the other (2063-1,-2 and 2053-2) (Figure S11B). This analysis suggests that by day 17, these two groups of hiPSC lines have accessed dorsal or ventral telencephalic fates with different efficiencies.

To gain a higher resolution view of transcriptional change in this system we decomposed the 6 hiPSC lines RNA-seq data with the informatic tool GWCoGAPS, identifying a set of 30 gene expression patterns (GWCoGAPS-III p1-p30) (Figure S12). Among these, patterns of genes restricted to the two groups of cell lines were identified. GWCoGAPS-III pattern p3, p14 and p15 showed differential expression in the two groups of cell lines at days 17 and 30 (Figure 5C and S12), again involving dorsal versus ventral telencephalic genes as indicated by projection of macaque developing cortex gene expression data set (Bakken et al., 2016) into GWCoGAPS-III using projectR (Figure 5Ci and ii). The GWCoGAPS-III p3 revealed enhanced expression of genes promoting dorsal telencephalic fates, including FEZF2 and PAX6 (p=5.0e-11 and p=9.0e-13 in DESeq2 on day 17). In contrast, the top weighted genes in the GWCoGAPS-III p15 included regulators of inhibitory neuron differentiation and function (ASCL1*, NKX2-1*, LHX6, LHX8, GAD1, DLX1*, DLX2*, DLX5*, DLX6*; *, p=1.0e-5 in DESeq2 analysis of differential expression on day 30) (Figure S11C).

These data raise the question whether the differential bias toward dorsal or ventral telencephalon shown by these hiPSC lines emerge from an earlier fate segregation process. GWCoGAPS-III p2 and p16 distinguished the two groups of cell lines already at day 8 (Figure 5Di). Interestingly, the projection of macaque developing cortex gene expression data (Bakken et al., 2016) into the GWCoGAPS-III patterns indicates that cortical hem genes, including OTX2, WNT8B, RSPO2, and WLS were highly weighted in p2 more than in p16 (Figure 5Di and S11D). These dynamics were confirmed by the high expression of LMX1A at day 8 followed by induction of EGFR at later time points in the lines with dorsal bias, consistent with the appearance of cortical excitatory neuronal precursors following patterning states (Figure 5Dii). These data indicate that the three lines that most efficiently generate cortical excitatory neurons were biased towards dorso-caudal organizer fates at earlier steps. The generality of the emergence of early organizer states *in vitro* was confirmed in public sequencing data of differentiating cortical neurons derived from hiPSCs (Edri et al., 2015; van de Leemput et al., 2014) by projection into the hem-associated GWCoGAPS-III p2 (Figure S11E). Importantly, NKX2.1 and SHH, well known ventral fate regulators (Sandberg et al., 2016), were maximally expressed in the cell lines with ventral bias on days 17 and 30 (Figure 5Dii and S11F). Prior to this, on day 8, FGF8 known to pattern the antero-ventral telencephalon (Storm et al., 2006) was maximally expressed in the cell lines with ventral bias (Figure 5Dii). These data indicate that the divergent neuronal trajectory bias of hNSCs was coordinated with early mechanisms regulating the emergence of distinct organizer signals in the generation of forebrain NSCs from pluripotent states (Figure 6). The molecular and cellular basis for this variation may be readily probed further across many human pluripotent lines, progressing *in vitro* as we describe here.

**Figure 6.**
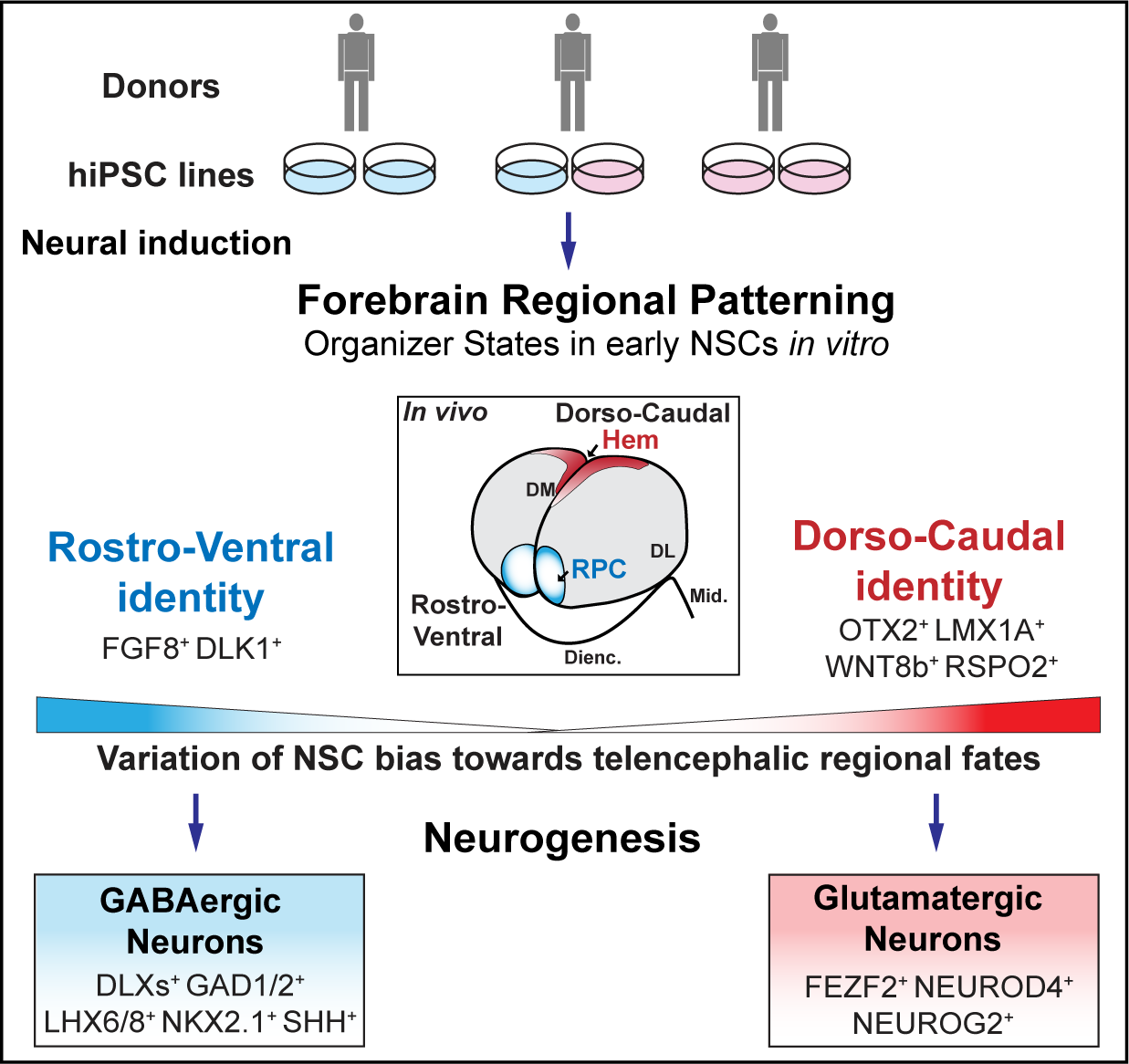
Divergent trajectory bias of human NSCs generated from pluripotent states. Variation in the emergence of organizer states during the generation of forebrain NSCs from donor specific hiPSCs, shown by different colors of the plates. The *in vivo* model illustrates developmental axes and signaling centers of the telencephalon. *In vitro,* early hNSCs show bias in their regional patterning state and progress from rostro-ventral or dorso-caudal identity towards inhibitory or excitatory neurogenesis. DM/DL: dorso-medial/-lateral.

## DISCUSSION

In the present study, we first defined the transcriptional progression of mouse and human cortical NSCs transitioning to glutamatergic excitatory neuronal fates *in vitro*. Our understanding of the cellular dynamics across dorsal telencephalic neuronal commitment was extended by the analysis of mouse NSC populations transiently expressing distinct levels of EGFR or PDGFR. We show that FGF2 signaling generates asymmetry within NSC progenitors instructing a wave of EGFR_high_ cells that take on cortical excitatory neuronal fates. Mouse and human EGFR_high_ NSCs in low doses of FGF2 initiated an endogenous BMP signaling cascade as cortical glutamatergic neurons first differentiated. In higher FGF2 concentrations we observed compromised neurogenesis that we attribute to a delayed BMP activity and differentiation of EGFR_high_ cells, preferentially generating glia fates in these conditions. The different neuronal trajectories traversed by *in vitro* NSCs exposed to varying FGF2 doses are consistent with previous *in vivo* results indicating that FGF2 signaling perturbation, during mouse embryogenesis, affects neurogenesis in the dorsal telencephalon specifically (Korada et al., 2002; Raballo et al., 2000; Rash et al., 2011; Rash et al., 2013). It has been shown in mouse that PDGFR*α* marks dorsal telencephalic NSCs that differentiate into neurons *in vivo* (Andrae et al., 2001) and *in vitro* after exposure to PDGF ligands (Johe et al., 1996; Park et al., 1999). We showed that PDGFR*α*_high_ NSCs do not respond to FGF2-induced BMP and generate GABAergic inhibitory more efficiently than cortical excitatory neurons in our *in vitro* system. A subpallial origin of these NSCs seems unlikely, as fluorescence activated cell sorting data in our lab, not shown in this work, indicate that the discrete PDGFRα_high_ and EGFR_high_ subtypes become one population double positive for both receptors in suspension, suggesting these are two states of the same precursor that probably reset potency with passage, as previously reported for NSCs (Ravin et al., 2008). It will be interesting to determine if these embryonic NSCs share features with other previously identified adult mouse telencephalic precursors expressing PDGFRα, including a subset of B type NSCs in the SVZ generating olfactory bulb interneurons and oligodendrocytes (Jackson et al., 2006), or others defining the oligodendrocyte lineage (Marques et al., 2016). Extending previous work identifying EGFR expressing cells as astro-glial precursors of the late SVZ in rodents and monkey (Burrows et al., 1997; Lillien and Raphael, 2000; Rash et al., 2019; Sun et al., 2005), our work defines a transient population of neurogenic EGFR_high_ RGCs responsive to BMP signaling undergoing acute neuronal commitment during cortical development *in vitro*.

From the first generation of neurons from mouse pluripotent sources *in vitro* (Kim et al., 2002; Okabe et al., 1996) to the many recent studies of the transcriptomic and epigenetic landscapes of the developing human cerebral cortex (de la Torre-Ubieta et al., 2018; Nowakowski et al., 2017; Zhu et al., 2018), it has been shown that neuronal differentiation from NSCs requires transitions through a series of cellular intermediates. Here, after defining the acute transition events leading to post-mitotic excitatory neuron fates, we explored *in vitro* the intrinsic patterning mechanisms that coordinate corticogenesis prior to the ingrowth of thalamic afferents *in vivo* (Armentano et al., 2007; Bishop et al., 2000; Cholfin and Rubenstein, 2007; Fukuchi-Shimogori and Grove, 2001; Miyashita-Lin et al., 1999; Nakagawa et al., 1999; Shimogori and Grove, 2005). We demonstrate the sequential appearance of organizer and cortical neuron precursor domains *in vitro*. Previous work has demonstrated the presence of organizer structures in cerebral organoids derived from human iPSCs (Renner et al., 2017). We show that the earliest events in human telencephalic patterning and neurogenesis may be systematically analyzed with PSCs using this 2D *in vitro* system, which give different advantages compared to 3D cultures (Pasca, 2018), and where we can observe the emergence of competing cortical patterning signals that determine regional neuronal fates.

The variation in the organizer states and subsequent divergent telencephalic trajectories revealed in the newly-generated iPSC lines validates this concept, providing an opportunity to define how morphogenetic spatial patterning is co-ordinated with the specification of excitatory or inhibitory neurons in the human telencephalon. Consistent with these data, another recent study shows variation in dorso-ventral telencephalic fate bias across cerebral organoids derived from multiple donor iPSCs (Kanton et al., 2019). Interestingly, we also observed that replicate lines from same individual (2053-2, −6) can traverse divergent telencephalic regional fates under same differentiation conditions. Previous reports have shown that transcriptional heterogeneity or differentiation capacity variability of multiple donor iPSC lines are under both genetic and epigenetic control (Carcamo-Orive et al., 2017; Nishizawa et al., 2016). Future studies will link genetic and epigenetic mechanisms to the divergent transcriptional phenotypes and fate bias we observed here.

The developmental mechanisms controlling these early telencephalic fates are of great interest as they are proximal to genetic risk for many neurodevelopmental disorders, including autism spectrum disorder (ASD) (Consortium, 2017; Kwan et al., 2012a; Madison et al., 2015; Marchetto et al., 2016; Mariani et al., 2015; Schafer et al., 2019), and brain cancers (Crawley et al., 2016; Ernst, 2016). Progress continues in defining disease relevant *in vitro* phenotypes using genetically distinct hPSC lines as a central tool in the development of novel therapeutic interventions (Fujimori et al., 2018; Hubler et al., 2018; Lang et al., 2018). Our study suggests that to achieve accurate models of risk for neuro-psychiatric disease, it will be necessary to more powerfully assess the extent and the origin of the developmental variation in patient-specific iPSCs, as they progress through the non-linear transitions described here. Defining, as we described, cell state transitions and subsequent distinct neuronal differentiation trajectories will be central in selecting optimal lines for specific fates and designing cell assays that more efficiently reveal phenotypes of interest, across a more precisely controlled neurogenic landscape *in vitro*. When integrated with the unprecedented transcriptomic and epigenetic mapping of the human forebrain (Amiri et al., 2018; de la Torre-Ubieta et al., 2018; Li et al., 2018; Wang et al., 2018; Zhu et al., 2018), the neural cell state transitions defined in this study will yield new functional insight into the origin of developmental risk for neuro-psychiatric disorders.

## AUTHOR CONTRIBUTIONS

N.M. and R.D.M. conceived the study. N.M. performed experiments and data collection. N.M. and SK.K. performed stem cell culture. M.D.B. performed e-physiology. Y.W. generated hiPSC lines. JH.S. generated bulk RNA-seq data. G.SO., E.J.F., JH.S., C.C. processed and analyzed bulk RNA-seq data. N.M. and J.A. collected monkey V1 single cells. S.M. and C.C. analyzed scRNA-seq data. N.M., SK.K., S.S., D.J.H. performed IF, imaging and high-content image analysis. N.M., C.C., G.SO, D.J.H. performed lineage analysis. N.M. and K.R.M. performed neuro-morphology analysis. B.G.R. performed IHC and imaging of the monkey tissue slides. J.A. banked monkey tissue. N.M., SK.K., G.SO., S.S., Y.W., N.A.O., J.G.C., C.C., D.J.H. developed biological and bio-informatic pipelines to manipulate and analyze hPSCs. N.M., SK.K., G.S.O., D.J.H., C.C., N.S., P.R., R.D.M. interpreted the data. A.J.C., R.B., N.J.B., D.R.W., N.S., P.R. and R.D.M. directed the research. N.M., SK.K., C.C., R.D.M. wrote the manuscript. All authors participated in discussion of results and manuscript editing.

## Supporting information

Micali et al_Supplementary Figures & Legends

## ACKNOWLEDGEMENTS

This work was supported by the Lieber Institute for Brain Development (LIBD) and the NIH grants DA02399 and EY002593 to P.R. and MH106934, MH106874, MH116488, MH110926 and MH109904 to N.S.. E.J.F. and G.S.O. were supported by the NIH (CA177669, CA212007, CA006973), the Johns Hopkins University Discovery Award, and the Johns Hopkins School of Medicine Synergy Award. We thank many members of LIBD and Yale Department of Neuroscience (Rakic and Sestan labs) for their helpful comments on this work and manuscript. We thank Dr. Alvaro Duque (Rakic Lab) and MacBrainResource (https://medicine.yale.edu/neurosci-ence/macbrain/) for providing NHP tissue (NIMH R01-MH113257 to A.D.).

