## Supplementary material for "Variation of human neural stem cells generating organizer states *in vitro* before committing to cortical excitatory or inhibitory neuronal fates": Micali et al_Supplementary Figures & Legends

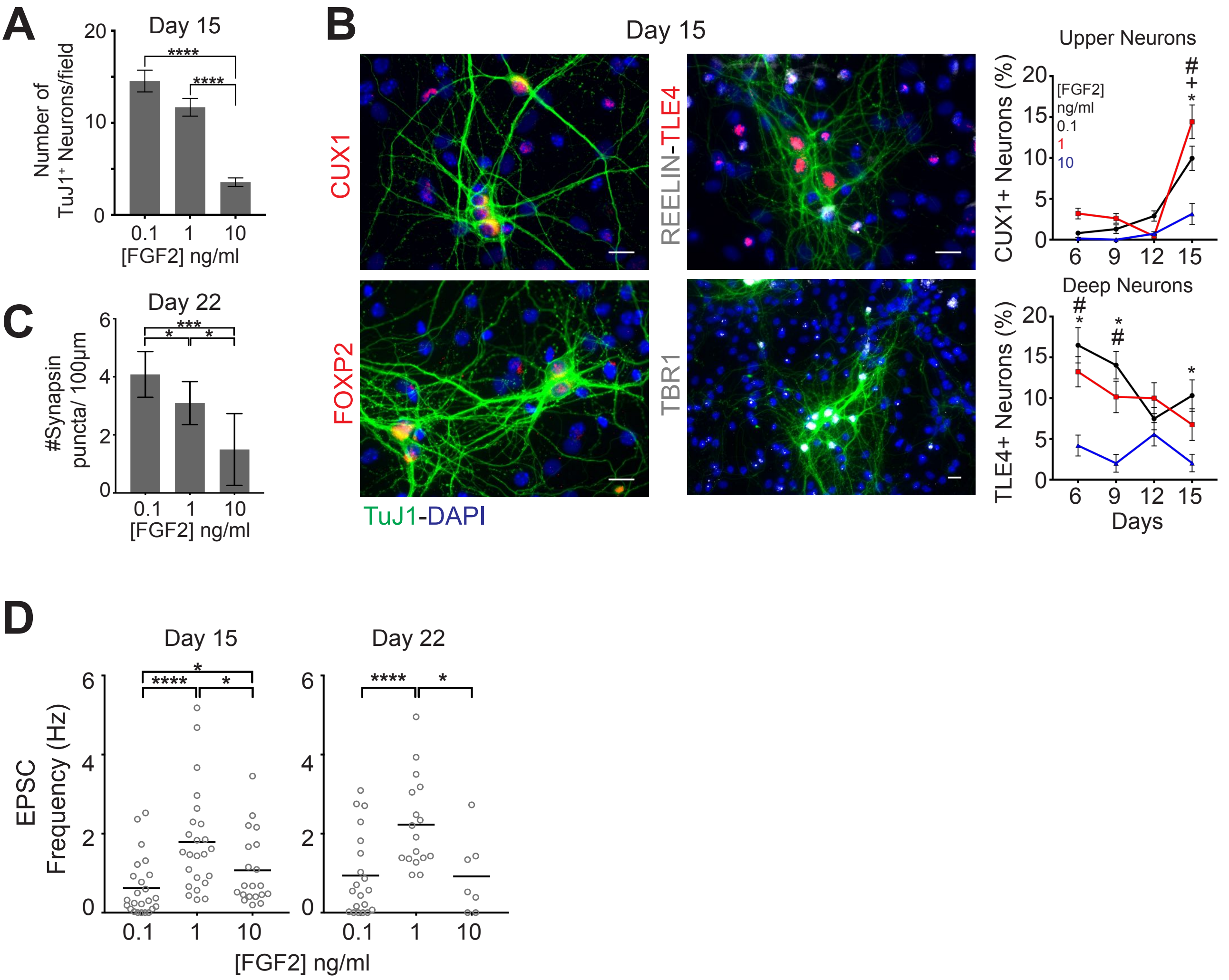


Figure S1-Related to Figure 1

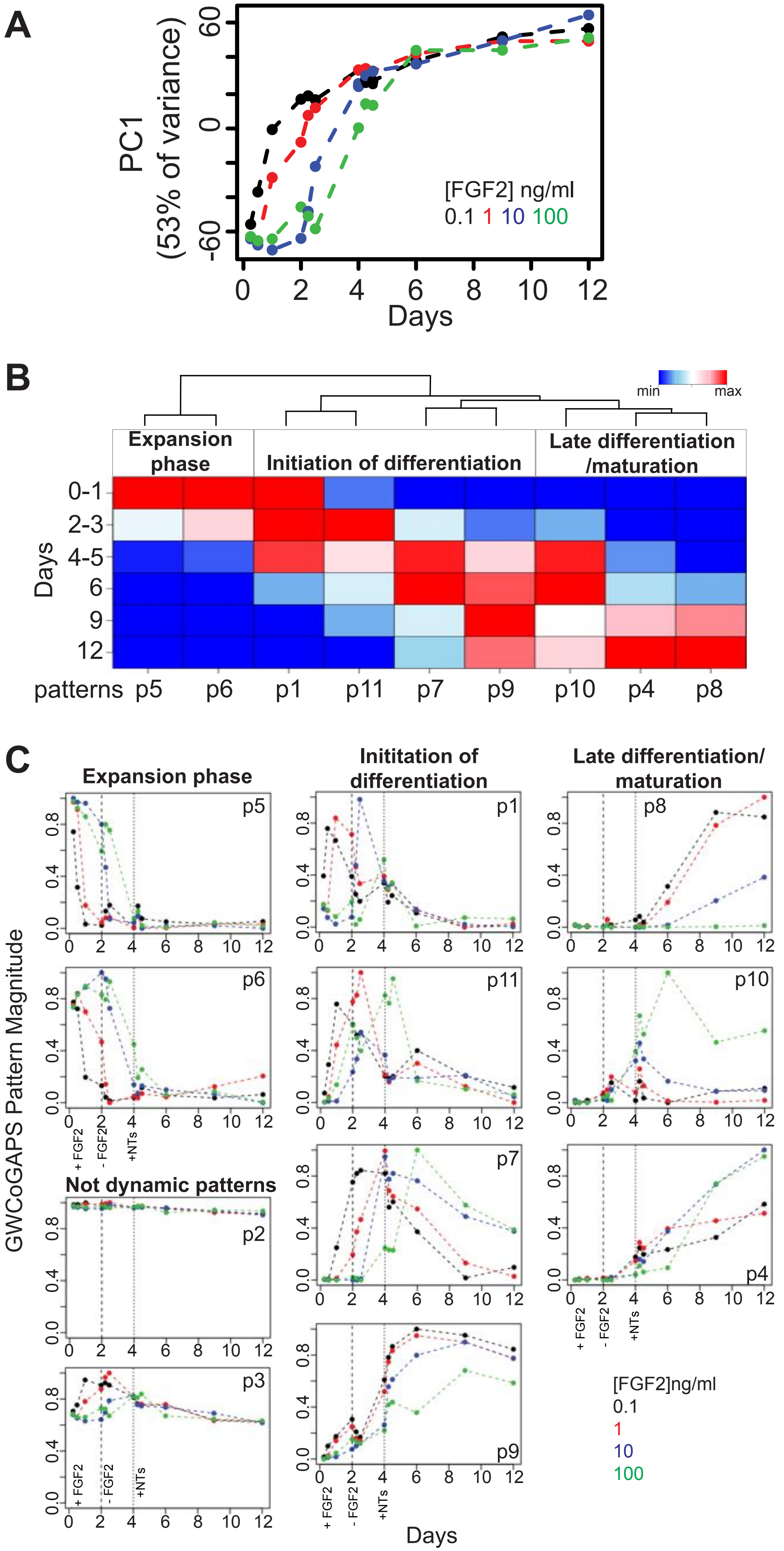


Figure S2-Related to Figure1

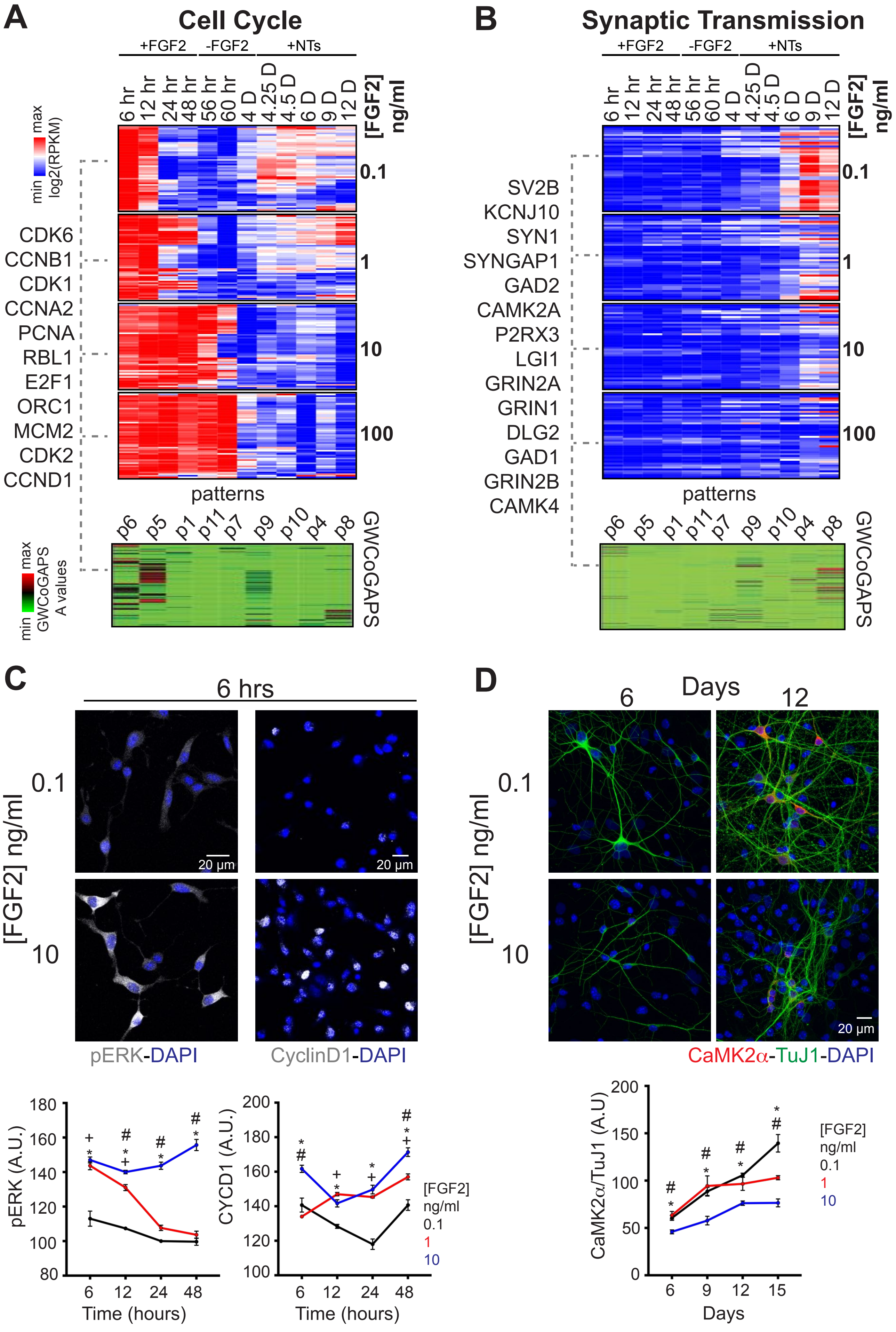


Figure S3-Related to Figure1

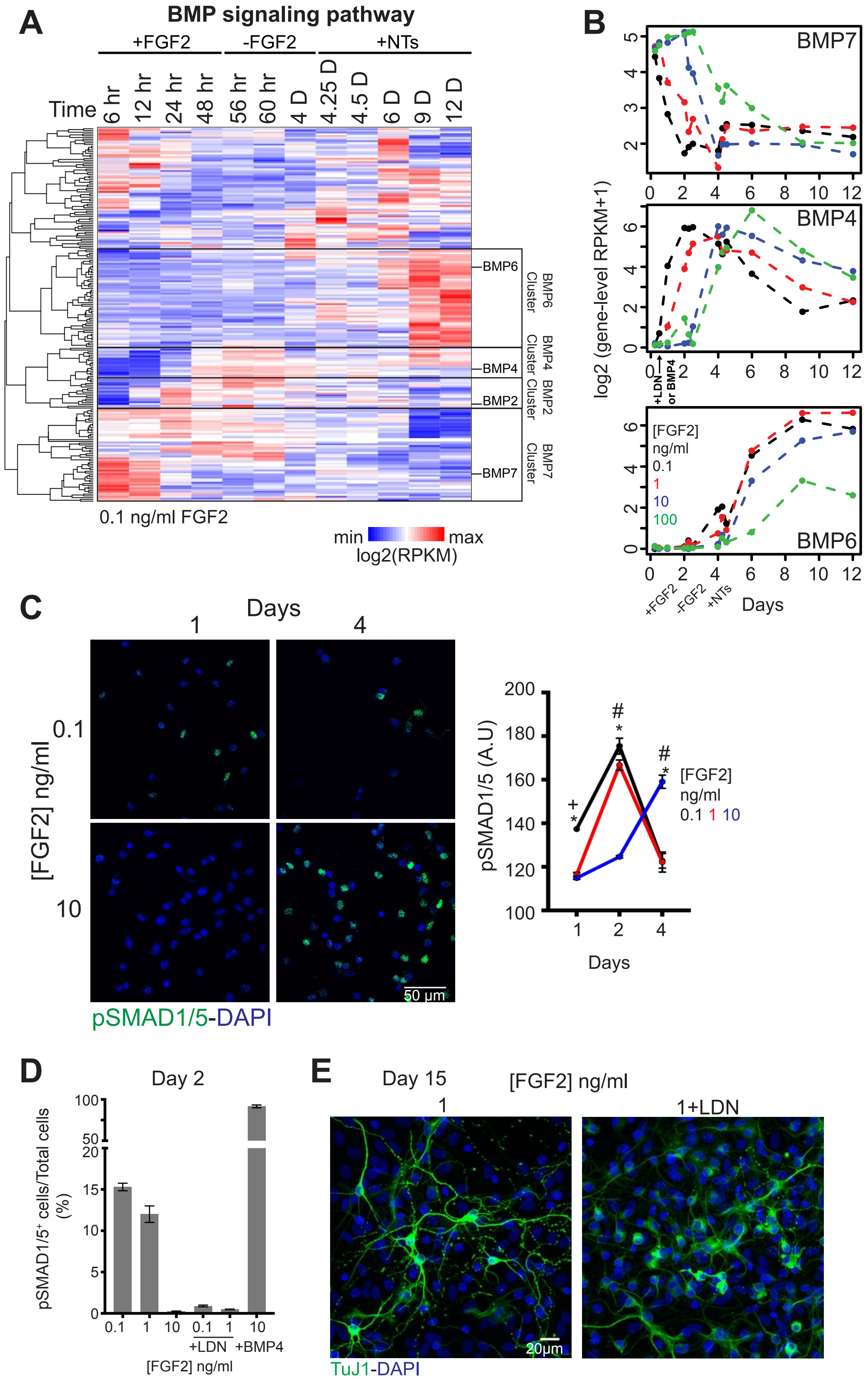


Figure S4-Related to Figure 1

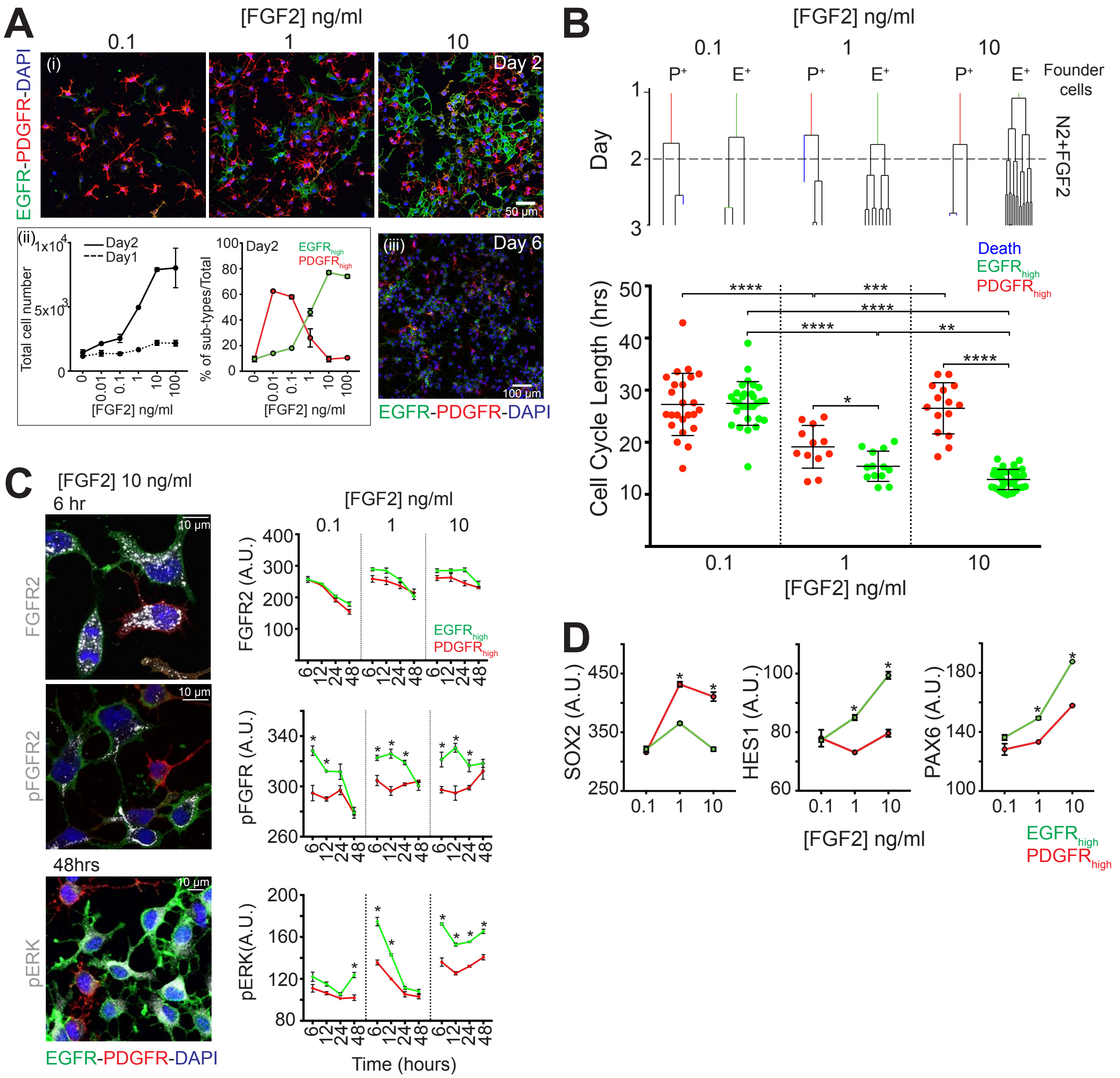


Figure S5-Related to Figure 2

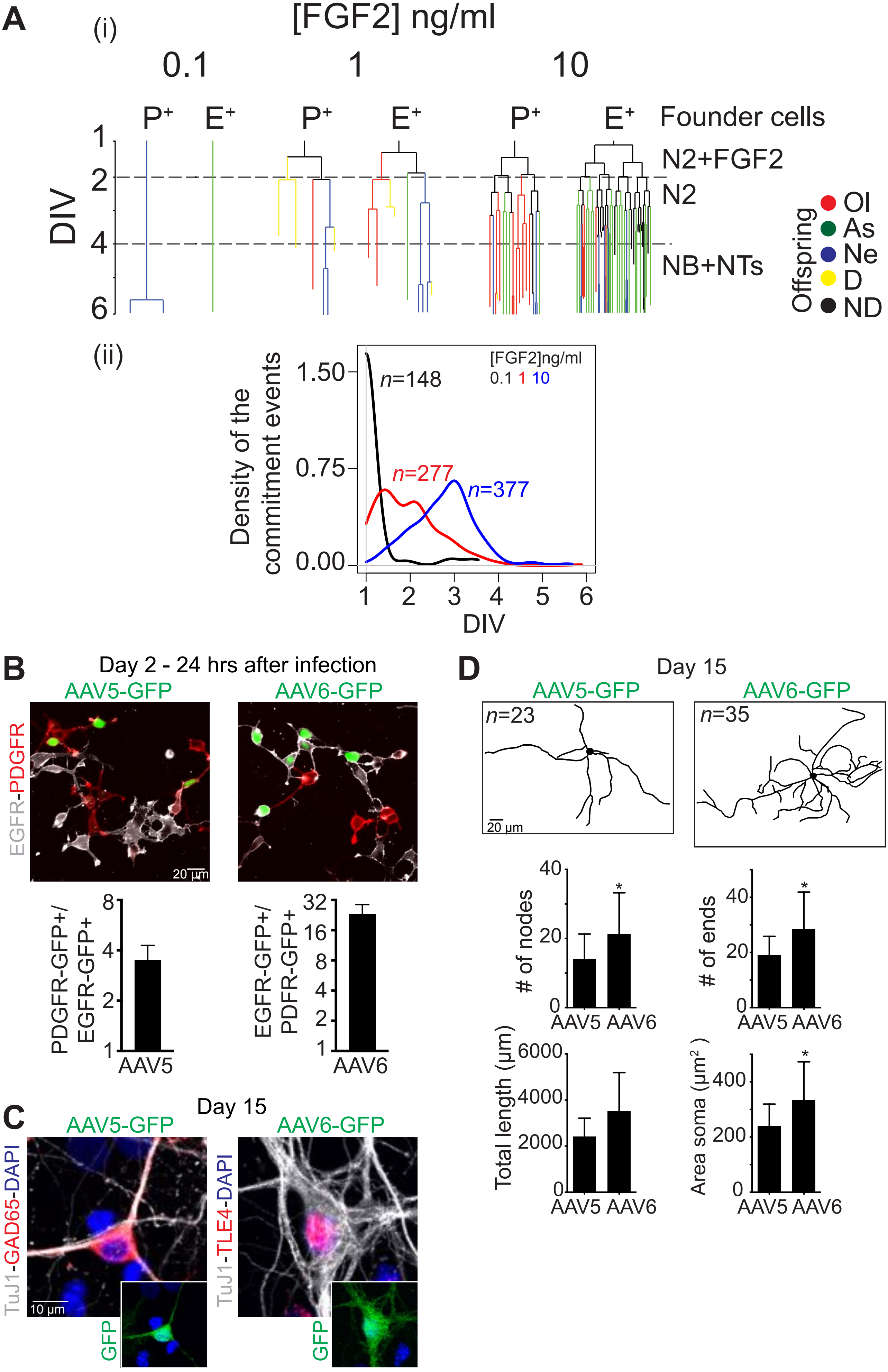


Figure S6-Related to Figure 2

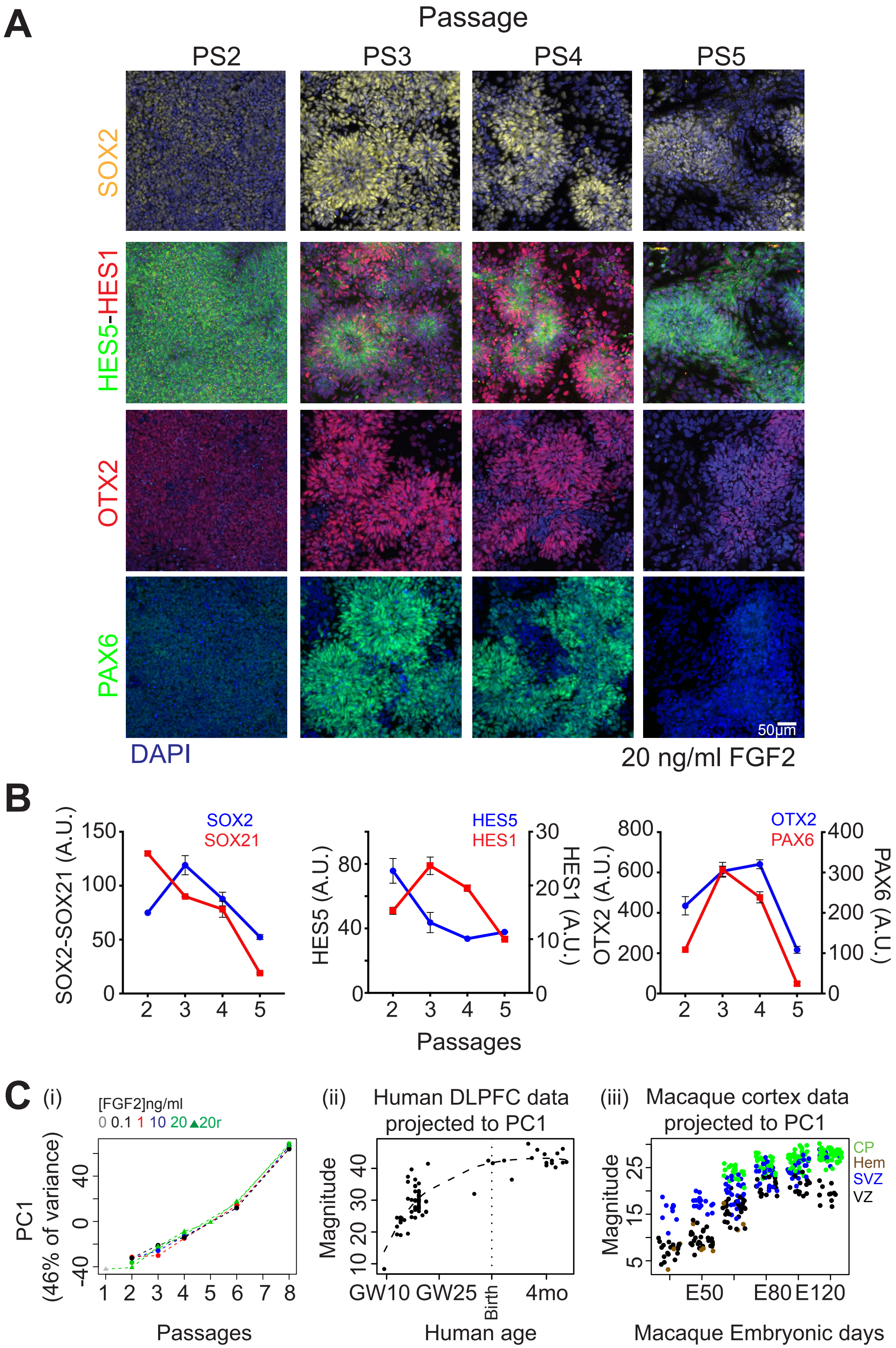


Figure S7-Related to Figure 3

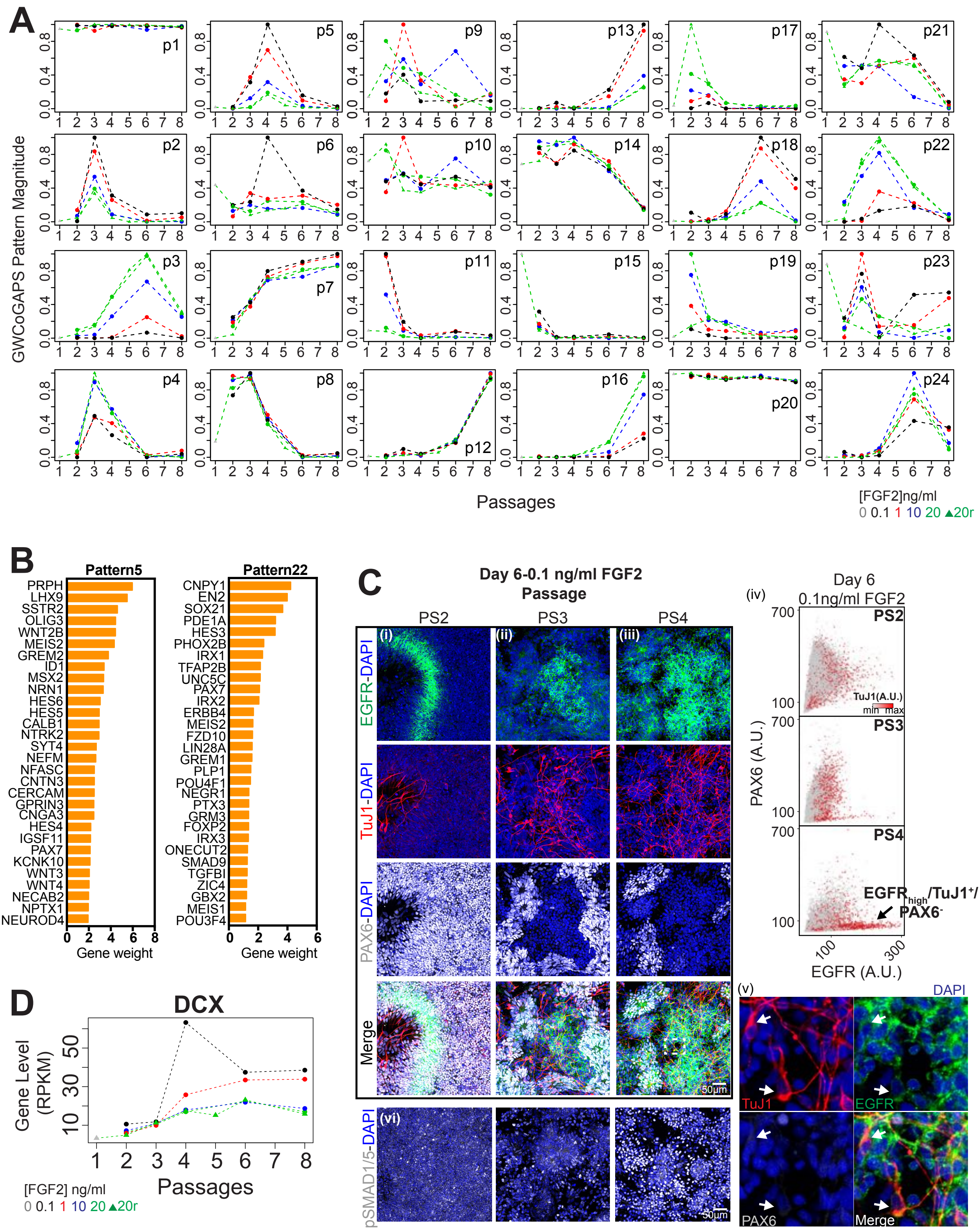


Figure S8-Related to Figure 3

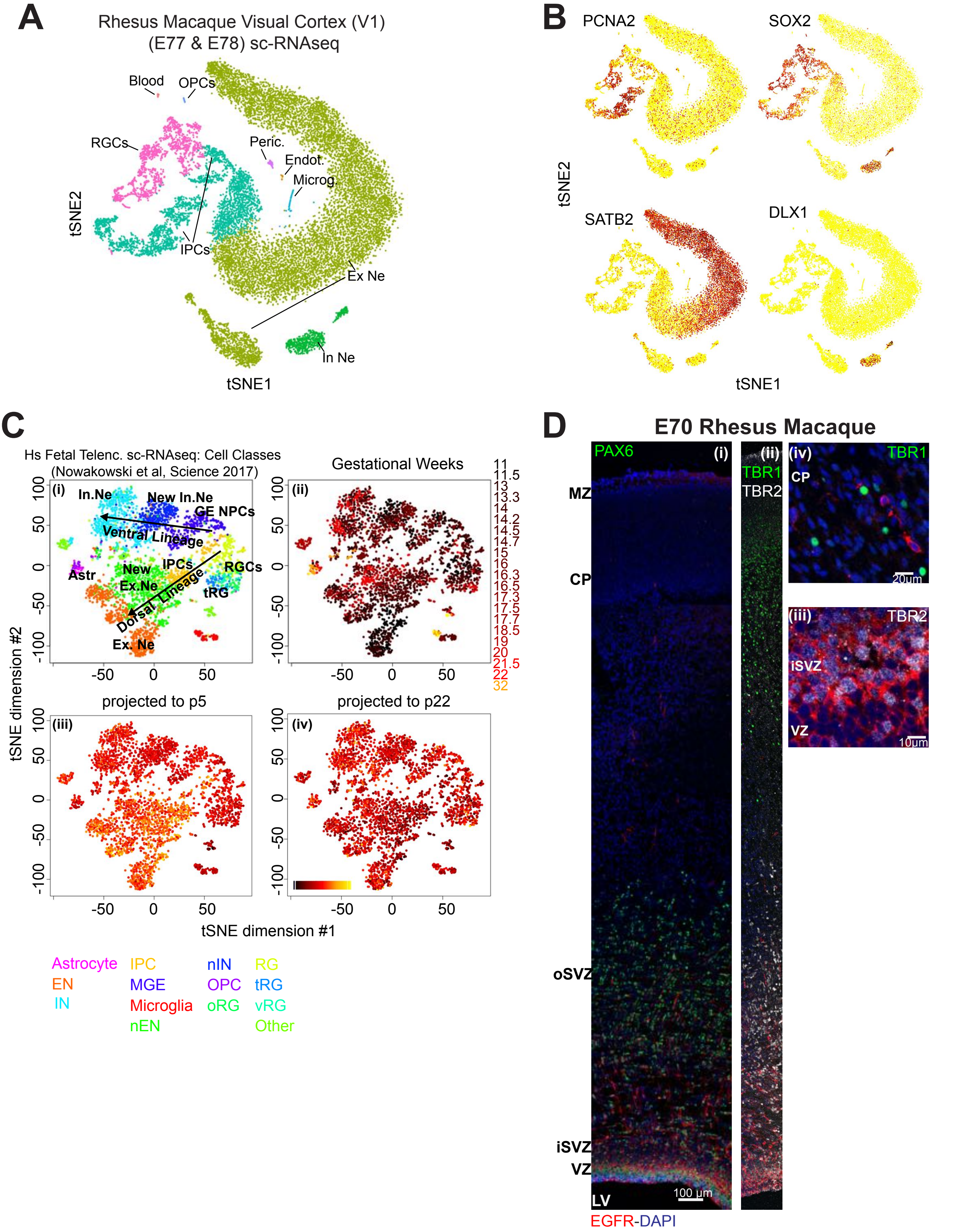


Figure S9-Related to Figure 3

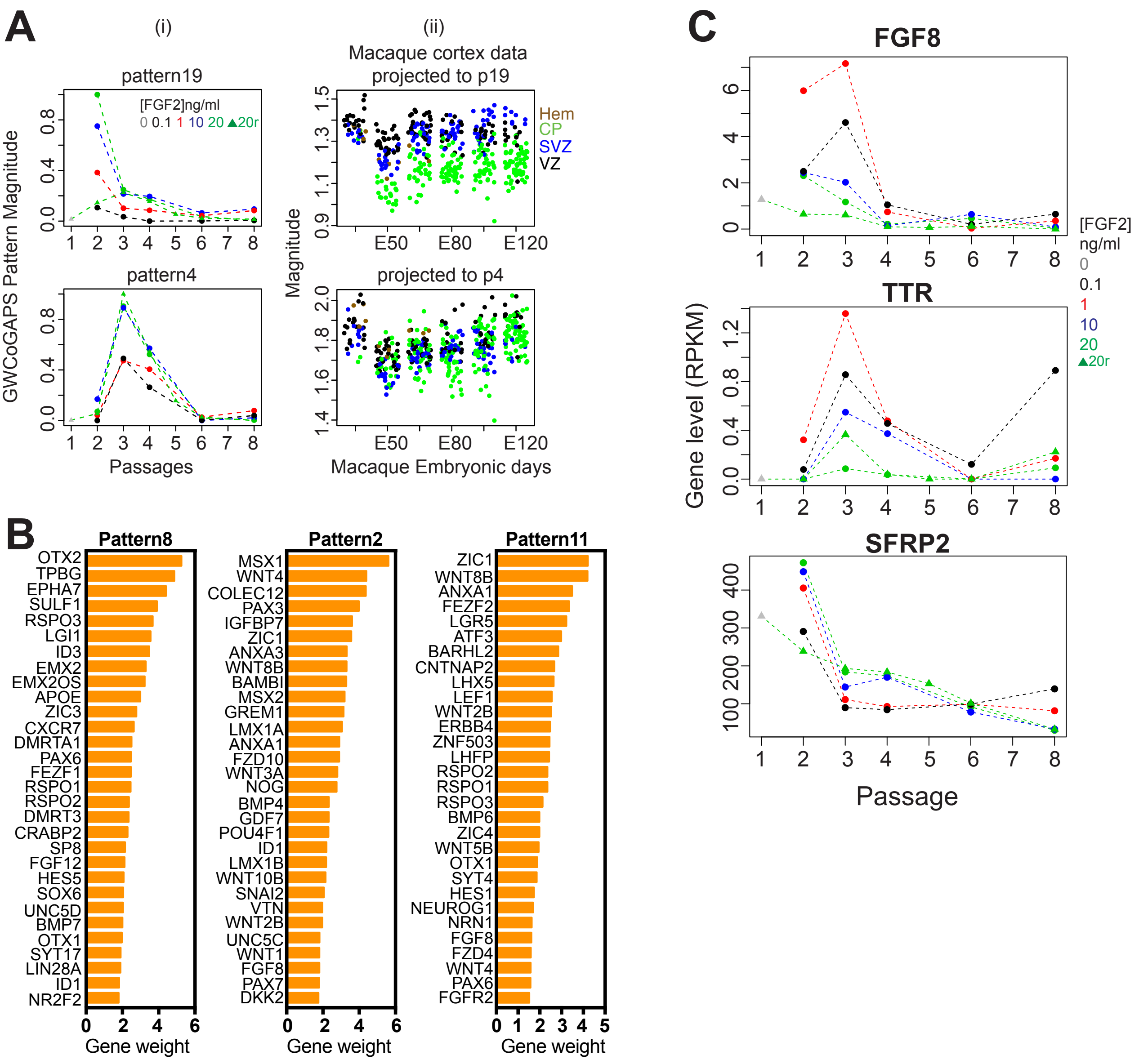


Figure S10-Related to Figure 4

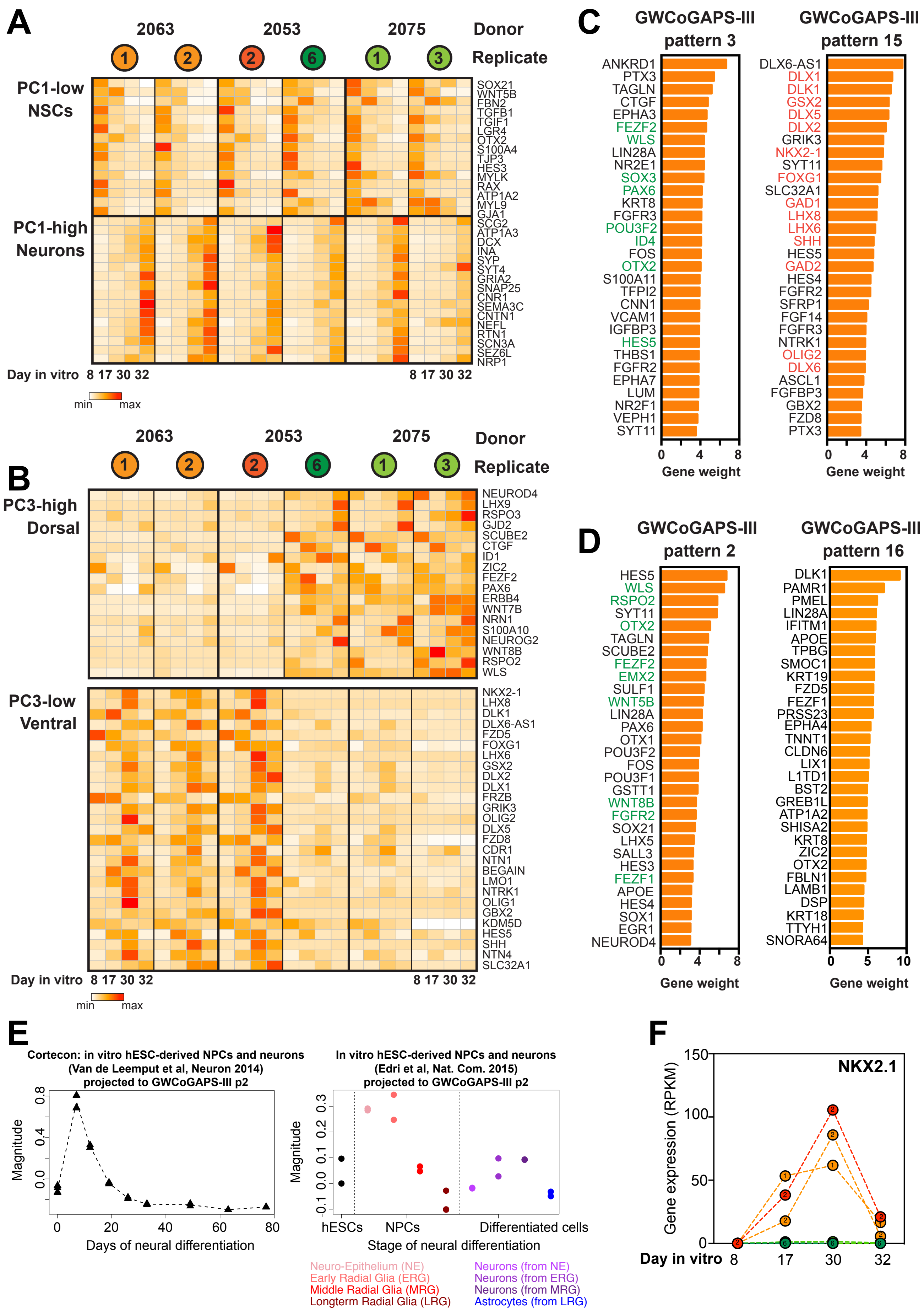


Figure S11-Related to Figure 5

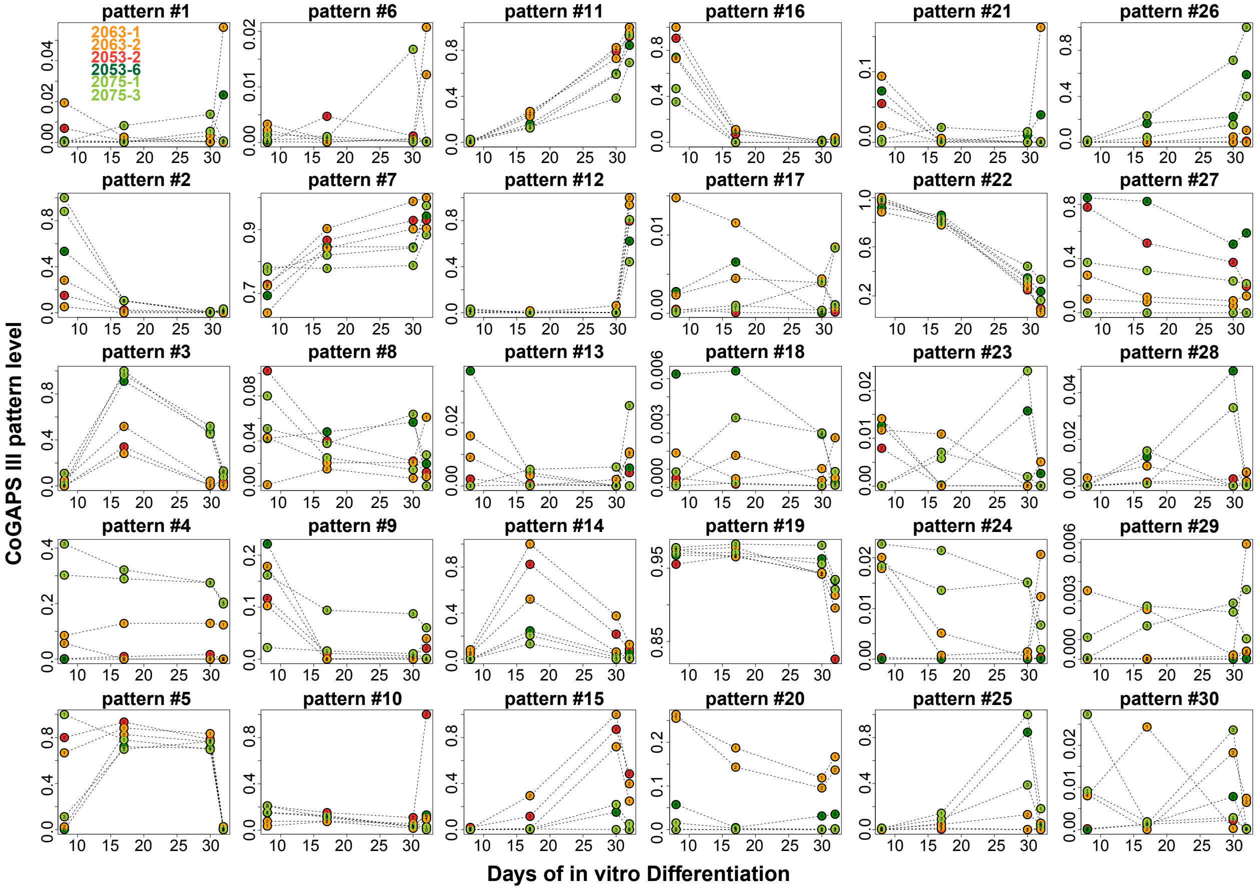


Figure S12-Related to Figure 5

### SUPPLEMENTARY FIGURE LEGENDS

**Figure S1-Related to Figure 1. Cortical neuron types and maturation regulated by early FGF2 signaling in mouse NSCs (A)** Mean number of $TuJ1^+$ neurons/field, from 4 independent DIV 15 cultures, t-test. (**B**) (Left) Immuno-fluorescence images of $TuJ1^+$ neurons, cultured over rat astrocytes, expressing CUX1, REELIN or TLE4, FOXP2, or TBR1 at DIV 15. Scale Bars, 20 µm. (Right) Percentage of $TuJ1^+$ neurons expressing TLE4 or CUX1. ANOVA comparing FGF2 doses ($p<0.05$). *: 0.1 vs 10. #: 1 vs 10. +: 0.1 vs 1. **(C)** Counts of individually segmented Synapsin1 puncta per 100 µm of neurite length also individually segmented in DIV 22 $TuJ1^+$ neurons. Mean + St.Dev., t-test n > 5 fields per measurement. **(D)** Mean frequency of Excitatory Post Synaptic Currents (EPSCs) in DIV 15 and 22 neurons derived from 0.1, 1 or 10 ng/ml FGF2 condition. t-test.

**Figure S2-Related to Figure 1. Transcriptional dynamics of *in vitro* mouse NSC differentiation** (**A**) PCA of gene-level RNA-seq data from 6 hours to 12 days of differentiation. NSCs were exposed to varying FGF2 doses and differentiated. (**B**) Hierarchical clustering of patterns defined by GWCoGAPS. (**C**) The 11 patterns defined by GWCoGAPS NMF algorithm across FGF2 doses and differentiation time, organized into three major groups, as defined by hierarchical clustering analysis in (B).

**Figure S3-Related to Figure 1. Cell cycle gene activity in NSCs related to neuronal differentiation** (**A**) (Top) Expression levels of selected GO “cell cycle” genes. Well know cell cycle genes are indicated on the left of the heat map. (Bottom) Gene weights for the same genes in each GWCoGAPS pattern. (**B**) (Top) Expression levels of selected GO “synaptic transmission” genes. Well know synaptic genes are indicated on the left of the heat map. (Bottom) Gene weights for the same genes in each GWCoGAPS pattern. Color key as in A. (**C**). (Top) Immuno-fluorescence images of pERK and Cyclin D1 expression in NSCs 6 hours after plating. Phosphorylation of MAPK/ERK and expression of Cyclin D1 were sensitive to FGF2 dose.

(Bottom) Mean signal intensity of pERK and Cyclin D1 (CYCD1) within individually segmented cells from high-throughput image analysis, across FGF2 doses and time, ± St.Dev. ANOVA comparing FGF2 doses ($p<0.05$). *: 0.1 versus 10; #: 1 versus 10; +: 0.1 versus 1. (**D**) Synaptic maturation genes increased when NSCs abruptly left the cell cycle in low FGF2. (Top) Immuno-fluorescence images of CaMK2α expression in TuJ1$^+$ neurons cultured over rat astrocytes, at DIV 6 and 12. (Bottom) Mean signal intensity of CaMK2α normalized over TuJ1 intensity, across FGF2 doses and time, ± St.Dev. ANOVA comparing FGF2 doses ($p<0.05$). *: 0.1 versus 10. #: 1 versus 10; +: 0.1 versus 1 FGF2. In low FGF2, NSCs down-regulate cell cycle genes and traverse later steps of neuron differentiation more efficiently than in high FGF2.

**Figure S4-Related to Figure 1. Endogenous BMP signaling dynamics across mouse *in vitro* neurogenesis** (**A**) Expression of genes in the "BMP receptor signaling" gene set from NCI (http://software.broadinstitute.org/gsea/msigdb/geneset_page.jsp?geneSetName=PID_BMP_PATHWAY). (In boxes) Gene clusters defined by hierarchical clustering containing different BMP ligands. 0.1 ng/ml FGF2 is shown. Genes correlated with BMP7, BMP4, BMP6 clusters were summarized in Figure 1. (**B**) Expression of BMP7, BMP4 and BMP6 from RNA-seq data. BMP signaling modulation with LDN and BMP4 is illustrated for BMP4 plot (**C**) (Left) Immuno-fluorescence images of pSMAD1/5 at Day 1 and 4 after NSC plating. (Right) Mean signal intensity of pSMAD1/5 within individually nuclear segmented cells from high-throughput image analysis, at the indicated days after NSC plating, ± St.Dev. ANOVA comparing FGF2 doses ($p<0.05$). *: 0.1 vs 10. #: 1 vs 10. +: 0.1 vs 1. (**D**) Proportion of individually segmented pSMAD1/5$^+$ NSCs from high-throughput image analysis, relative to total cell number at DIV 2 for 0.1, 1 or 10 ng/ml FGF2; 0.1 and 1 ng/ml FGF2 plus LDN193189 (100 nM); 10 ng/ml FGF2 plus BMP4 (10 ng/ml); LDN or BMP4 were added 12 hours after plating. Mean values + St.Dev. (**E**) Immuno-fluorescence images of DIV 15 TuJ1$^+$ neurons, over rat astrocytes, derived from 1 ng/ml FGF2 condition plus or minus LDN (100 nM), added 12 hours after plating and withdrawn with FGF2 at DIV 2. The image shows well differentiated neurons with defined dendrites in control vs compromised TuJ1$^+$ cells with aberrant neuron morphology derived from NSCs exposed to LDN.

**Figure S5-Related to Figure 2. Selective response to FGF2 signaling by mouse NSC subtypes** (**A**) (i) Immuno-fluorescence images of PDGFR$\alpha$ and EGFR expression in NSCs at day 2 of FGF2 modulation; (ii) (Left) Proliferation curves of NSCs at day 1 and day 2 after plating. (Right) Proportion of PDGFR$\alpha_{high}$ and EGFR$_{high}$ cells at day 2 after plating, from the high-throughput density plot shown in Figure 2A; (iii) Images of PDGFR$\alpha$ and EGFR expression in differentiated cells at DIV 6, after exposure to 10 ng/ml FGF2 in the first 2 days. (**B**) (Top) Lineage dendrograms from 2 days time-lapse recording. Colored segments reflect fluorescence signal at time T=0 for PDGFR$\alpha_{high}$ (red) vs EGFR$_{high}$ (green) founder cells. Recording started at DIV 1, T=0. Death fated cells are indicated in blue. (Bottom) Duration of the first inter-mitosis intervals. Mean length ± St.Dev. t-test. (**C**) (Left) Immuno-fluorescence images of EGFR$_{high}$ and PDGFR$\alpha_{high}$ NSCs expressing FGFR2 and pFGFR at 6 hours, and pERK at 48 hours after plating in 10 ng/ml FGF2. Scale bars are indicated. (Right) Mean signal intensity of FGFR2, pFGFR and pERK level within individually segmented EGFR$_{high}$ and PDGFR$\alpha_{high}$ cells, from high-throughput image analysis, ± St.Dev. t-test. (**D**) Mean signal intensity of SOX2, HES1 and PAX6 expression within individually nuclear segmented EGFR$_{high}$ and PDGFR$\alpha_{high}$ cells, from high-throughput image analysis, ± St.Dev. t-test.

**Figure S6-Related to Figure 2. Distinct fate bias of EGFR$_{high}$ and PDGFR$\alpha_{high}$ NSCs** (**A**) (i) Representative lineage dendrograms from 5 days time-lapse recordings of PDGFR$\alpha_{high}$ and EGFR$_{high}$ founder NSCs cultured with different doses of FGF2 and identified at DIV 1, to derivative neurons, oligodendrocytes or astrocytes, identified by expression of TuJ1, O4 and GFAP at DIV 6 (see Video S1, 2, 3). Recording started at DIV 1, T=0 . Ol, oligodendrocyte; As, astrocyte; Ne, neuron; D, death (apoptotic); ND, not determined. (ii) Commitment events density plot derived from all the cells traced, independently of the receptor expression. The plot shows that faster cell cycle exit correlates with early fate commitment of the NSCs. Number of cells traced per condition: 1106 in 10 ng/ml FGF2; 646 in 1 ng/ml FGF2; 202 in 0.1 ng/ml FGF2. *n*= number of terminal offspring cells for each lineage analysis. (**B**) (Top) Immuno-fluorescence images of PDGFR$\alpha_{high}$ and EGFR$_{high}$ NSCs 2 days after plating and 24 hours after infection with AAV5-GFP or AAV6-GFP. (Bottom) Infection efficiency for AAV5 and AAV6 adeno-serotypes, determined respectively as rate of PDGFR$\alpha_{high}$ / EGFR$_{high}$ cells infected by AAV5 and EGFR$_{high}$ / PDGFR$\alpha_{high}$ cells infected by AAV6, 24 hours after infection. (**C**) Immuno-fluorescence images of AAV5-GFP or AAV6-GFP DIV 15 TuJ1$^{+}$ neurons expressing GAD65 or TLE4. (**D**) (Top) Morphological analysis of AAV5-GFP and AAV6-GFP neurons. *n*= number of neurons traced. (Bottom) Mean values of the parameters analyzed ± St.Dev. t-test.

**Figure S7-Related to Figure 3. Dorsal telencephalic NSCs generated from human PSCs *in vitro*** (**A**) Immuno-fluorescence of fate regulators SOX2, HES5 and HES1, OTX2, PAX6 expression in hNSCs from PS1 to PS5, at 6 days after passage. (**B**) Mean fluorescence intensity in individual segmented cells from high-throughput image analysis, for the indicated markers at every passage. ± St.Dev. (**C**) (i) PCA of gene-level RNAseq data from hNSCs. FGF2 doses are indicated. 0 ng/ml FGF2 refers to PS1; 20r refers to PS2-PS8 sample replicates for 20 ng/ml FGF2 (ii) Projections of human DLPFC RNAseq data (Jaffe et al., 2018) into PC1; (iii) Projection of laser micro-dissected macaque cortex microarray data (Bakken et al., 2016) into the PC1. PC1 described a progressive change *in vitro* that parallels the *in vivo* development of the cortex, as CP emerges from the germinal VZ and SVZ. CP, cortical plate; Hem, cortical Hem; SVZ, subventricular zone; VZ, ventricular zone;.

**Figure S8-Related to Figure 3. Cortical neuron differentiation bias of human NSCs varies across passages** (**A**) 24 patterns defined by GWCoGAPS. FGF2 doses are indicated. 0 ng/ml FGF2 refers to PS1; 20r refers to PS2-PS8 sample replicates for 20 ng/ml FGF2 (**B**) 30 selected genes among the top 100 most strongly contributing to patterns p5 and p22, shown in Figure 3B. Pattern p5 is enriched in genes highly expressed during neuronal differentiation and CP formation. Pattern p22 is enriched in genes highly expressed by neural stem cells. (**C**) (Left) Immuno-fluorescence of hNSCs grown in 0.1 ng/ml FGF2 for 6 days showing expression of EGFR, TuJ1 and PAX6 in PS2 (i), PS3 (ii) and PS4 (iii). (iv) Scatter plot of individually segmented

EGFR vs PAX6 expressing cells, colored by TuJ1 expression at PS2, PS3 and PS4, from high-throughput image analysis. (v) Higher magnification of the dashed area in (iii-merge) showing EGFR/TuJ1 co-expressing cells, indicated with arrows. (vi) Immuno-fluorescence images of hNSCs grown with 0.1 ng/ml FGF2 for 6 days showing higher induction of pSMAD1/5 in PS4. Nuclei are stained with DAPI. (**D**) Dose-response expression of Doublecortin (DCX) through passages, from hNSC RNAseq data. FGF2 doses are indicated. 0 ng/ml FGF2 refers to PS1; 20r refers to PS2-PS8 sample replicates for 20 ng/ml FGF2.

**Figure S9-Related to Figure 3. Neurogenic transition states of RGCs in the developing primate cortex** (**A**) t-distributed stochastic neighbor embedding (tSNE) plots colorored by cluster annotation from macaque E77 and E78 Visual Cortex (V1) single cell RNA-seq data. 17161 cells (9193 cells from E77 V1 and 7963 from E78 V1) were analyzed. RGCs: radial glial cells; IPCs: intermediate precursors cells; OPCs: oligodendrocytes precursor cells; Peric.: pericytes; Endot.: endothelial cells; Microg.: microglia; ExNe: excitatory neurons; InNe: interneurons (**B**) tSNE plots colorored by markers for the major cell clusters: neurons (SATB2); cyclin RGCs and IPCs (PCNA); RGCs and IPCs (SOX2); IPCs (Eomes/TBR2 and NEUROG1) see also Figure 3E; interneurons (DLX1). (**C**) (i-iv) tSNE plots of human fetal telencephalic single-cell RNA-seq data of 3495 cells from (Nowakowski et al., 2017). (i) Depiction of cell classes across dorsal and ventral telencephalic developmental trajectories. Cell clusters nomenclature at the bottom of the figure. (ii) Ages of the donors from which cells derived. (iii and iv) Projections of the human fetal telencephalic single-cell RNA-seq data from (Nowakowski et al., 2017) into p5 and p22, colored by magnitude of projection, confirming that p5 is correlated with dorsal post-mitotic neuron signature *in vivo*. MGE NPCs, medial ganglionic eminence neural precursors; IPCs, intermediate progenitor cells; RGCs, radial glial cells; EN: excitatory neurons; IN, inhibitory neurons; nEN, new excitatory neurons; nIN, new inhibitory neurons; OPC, oligodendrocyte progenitor cells; oRG, outer radial glial cells; RG, radial glial cells; tRG, truncated radial glial cells; vRG, ventricular radial glial cells. (**D**) (i-iv) Immuno-histochemistry of E70 rhesus macaque dorsal parietal cortex tissues sections for EGFR (red; all panels) together with Pax6 (i), TBR1 and TBR2 (ii), TBR2 in the VZ and iSVZ (iii), TBR1 in CP(iv). Nuclei stained with DAPI.

**Figure S10- Related to Figure 4. Organizer identities of early passage human NSCs** (**A**) (i) GWCoGAPS patterns p19 and p4. FGF2 doses are indicated. 0 ng/ml FGF2 refers to PS1; 20r refers to PS2-PS8 sample replicates for 20 ng/ml FGF2. (ii) Projections of microarray data from laser micro-dissected macaque cortex (Bakken et al., 2016) into p19 and p4. CP, cortical plate; SVZ, subventricular zone; VZ, ventricular zone. (**B**) 30 selected genes among the top 100 most strongly contributing to patterns p8, p2 and p11 shown in Figure 4, enriched in genes highly expressed in the cortical hem. (**C**) Dose-response expression of FGF8, TTR, SFRP2 through passages from RNAseq data. FGF2 doses are indicated. 0 ng/ml FGF2 refers to PS1. 20r refers to PS2-PS8 sample replicates for 20 ng/ml FGF2.

**Figure S11-Related to Figure 5. Human NSC line variation in organizer states results in differential neuronal fate trajectories** (**A**) Heatmap showing expression levels of NSC - and neuron- related genes that are highly represented at the extremes of PC1, demonstrating the transition from neural precursors to neurons in all 6 hiPSC lines. (**B**) Heatmap showing expression levels of dorsal and ventral fate specification-related genes that are highly represented at the extremes of PC3, demonstrating distinct regional biases of each group of lines. (**C**) 30 selected genes among the top 100 genes most strongly contributing to the GWCoGAPS-III p3 and p15 shown in Figure 5C. Well-known dorsal (green) and ventral (red) fate specification-related genes are highlighted. (**D**) 30 selected genes among top 100 genes most strongly contributing to the GWCoGAPS-III p2 and p16 shown in Figure 5D. Well-known genes of the cortical hem (in green) are highly expressed at day 8 in the lines with dorsal lineage bias. (**E**) Projection of sequencing data of differentiating cortical neurons derived from hiPSCs from (Edri et al., 2015; van de Leemput et al., 2014) into the hem-associated GWCoGAPS-III p2. The data confirm the emergence of early organizers in *in vitro* hNSCs generated with other protocols in other labs. NSC types classified in (Edri et al., 2015) at the bottom. (**F**) Expression of the ventral gene NKX2.1 across the neural differentiation of the 6 lines. The marker is expressed at late time points by the ventral biased hNSC lines.

**Figure S12-Related to Figure 5. Human NSC line variation in organizer states results in differential neuronal fate trajectories** (**A**) 30 patterns defined by GWCoGAPS of the 6 hiPSC lines data set, indicated as GWCoGAPS-III p1-p30.
